# Kinesin-3 KIF14 Regulates Intraflagellar Transport Dynamics in Primary Cilia

**DOI:** 10.1101/2025.03.20.644298

**Authors:** Erika Mikulenková, Petra Pejšková, Roman Podhájecký, Luděk Štěpánek, Martina Huranová, Vladimír Varga, Zdeněk Lanský, Lukáš Čajánek

## Abstract

The primary cilium is an antenna-like organelle that plays a crucial role in development and homeostasis. Its growth and functions depend on the transport of cargo from the cilia base to the tip by intraflagellar transport (IFT) proteins and kinesin-2 motors. Primary cilia exhibit great variation in morphology and function across cell types and organisms. This diversity is thought to be conferred by the modulation of IFT by additional factors. However, examples of such non-canonical regulators, such as kinesin motors distinct from kinesin-2, are sparse. Thus, the involvement of non-canonical ciliary kinesins in intraciliary transport is unclear.

Here, we show that a poorly characterized member of the kinesin-3 family, KIF14, plays a prominent role in primary cilia trafficking in human cells. Using live cell imaging and expansion microscopy, we demonstrate that KIF14 depletion leads to impaired IFT. Furthermore, using TIRF microscopy we show that the motility of KIF14 and its effects on cilia trafficking rely on the motor domain of KIF14, which drives the processive runs along the ciliary axoneme in cells and *in vitro*, in co-operation with the C-terminal CC1 domain. Finally, we demonstrate that C-terminal truncations of KIF14, including patient mutation Q1380x, disrupt IFT by causing traffic jam-like accumulations of ciliary components in the distal part of the primary cilia, leading to bulged tips of cilia.

In summary, our data exemplify the role of a non-canonical ciliary kinesin in the regulation of ciliary trafficking and suggest a new paradigm for how kinesin-related trafficking defects may contribute to the pathology of disorder with a ciliary contribution.

## Introduction

Primary cilia are hair-like structures found on the surface of most vertebrate cells. Unlike motile cilia, primary cilia are non-motile and were initially considered vestigial, leading to their relative neglect in early research. However, recent studies have demonstrated their crucial role in responding to various external stimuli. Primary cilia contain receptors and messenger molecules for multiple signaling pathways, playing an essential part in regulating embryonic development and maintaining tissue homeostasis (1,2). Mutations that impair ciliary function result in a range of genetic disorders collectively referred to as ciliopathies, including conditions such as Joubert Syndrome, Nephronophthisis, and Meckel-Gruber syndrome (3,4). Therefore, a thorough understanding of cilia biology is crucial for uncovering the pathological mechanisms underlying these cilia-related diseases and for developing effective treatments.

A fully grown primary cilium is composed of the basal body, a mother centriole anchored to the plasma membrane via its distal appendages; the transition zone, a specialized domain at the base sorting proteins entering the ciliary compartment; and the axoneme, a microtubule-based rod protruding into the extracellular space and enclosed within the ciliary membrane (5). The growth of the cilium is mediated by Intraflagellar Transport (IFT) – a bidirectional movement of multiprotein particles termed IFT trains (6) along the doublet microtubules from the base of the cilium to its tip (anterograde direction) and back from the tip to the base (retrograde direction) (7,8). IFT is indispensable for cilia formation and maintenance from protists to vertebrates (8,9). The available evidence puts the kinesin-2 family of motors as exclusive regulators of anterograde IFT in most cilia (10–14). These canonical kinesin motors include the kinesin-2 heterotrimer (composed of KIF3A, KIF3B, and the kinesin-associated protein KAP-1) and the KIF17 homodimer (9,15). Additional kinesin motors localize to primary cilia (15,16) (we refer to them as non-canonical ciliary kinesins), including kinesin-3 motors KIF13B and KIF14 (17–19). These non-canonical ciliary kinesins are hypothesized to act as accessory motors to cooperate with the canonical kinesin-2 motors to control axoneme assembly, dynamics, and length (15,20). As primary cilia of individual cell types show structural and functional customizations (21), their diversification is proposed to be conferred by the activities of the non-canonical ciliary kinesin motors. Indeed, KLP-6 (an ortholog of KIF13B in *C. elegans*) regulates IFT in specialized primary cilia in male worms (19). However, evidence of non-canonical ciliary kinesins modulating IFT in vertebrate primary cilia is lacking.

KIF14, a member of the kinesin-3 family, is a plus-end directed motor, with an atypical disordered region found N-terminally to its motor domain. The disordered region and protein dimerization via C-terminal regions are implicated in the regulation of KIF14 processive movement *in vitro* (22). KIF14 has been linked to the regulation of chromosome segregation and cytokinesis (23–26). Our previous work established KIF14 as a ciliary component and regulator of primary cilia formation in human cells (17). In accordance with that, KIF14 has been implicated in brain development, with KIF14 mutations being described in patients with primary microcephaly or Mecker-Gruber syndrome (27–29). Here, we demonstrate that KIF14 moves processively along the ciliary axoneme, modulates IFT, and its C-terminal truncated mutants impair IFT due to traffic-jam-like protein assemblies. Our data provide the first evidence of IFT fine-tuning by non-canonical ciliary kinesin in primary cilia of vertebrate cells.

## Results and discussion

### KIF14 is required for correct IFT in primary cilia

To reveal the function of KIF14 in primary cilia, we used telomerase-immortalized retinal pigment epithelial hTERT-RPE-1 cells, where ciliogenesis can be readily triggered by serum deprivation. The general outline of our experiment is depicted in Fig. 1A. Following siRNA transfection, we first confirmed that KIF14 siRNA efficiently depletes KIF14 protein levels (Suppl. Fig. 1A).

**Figure 1.**
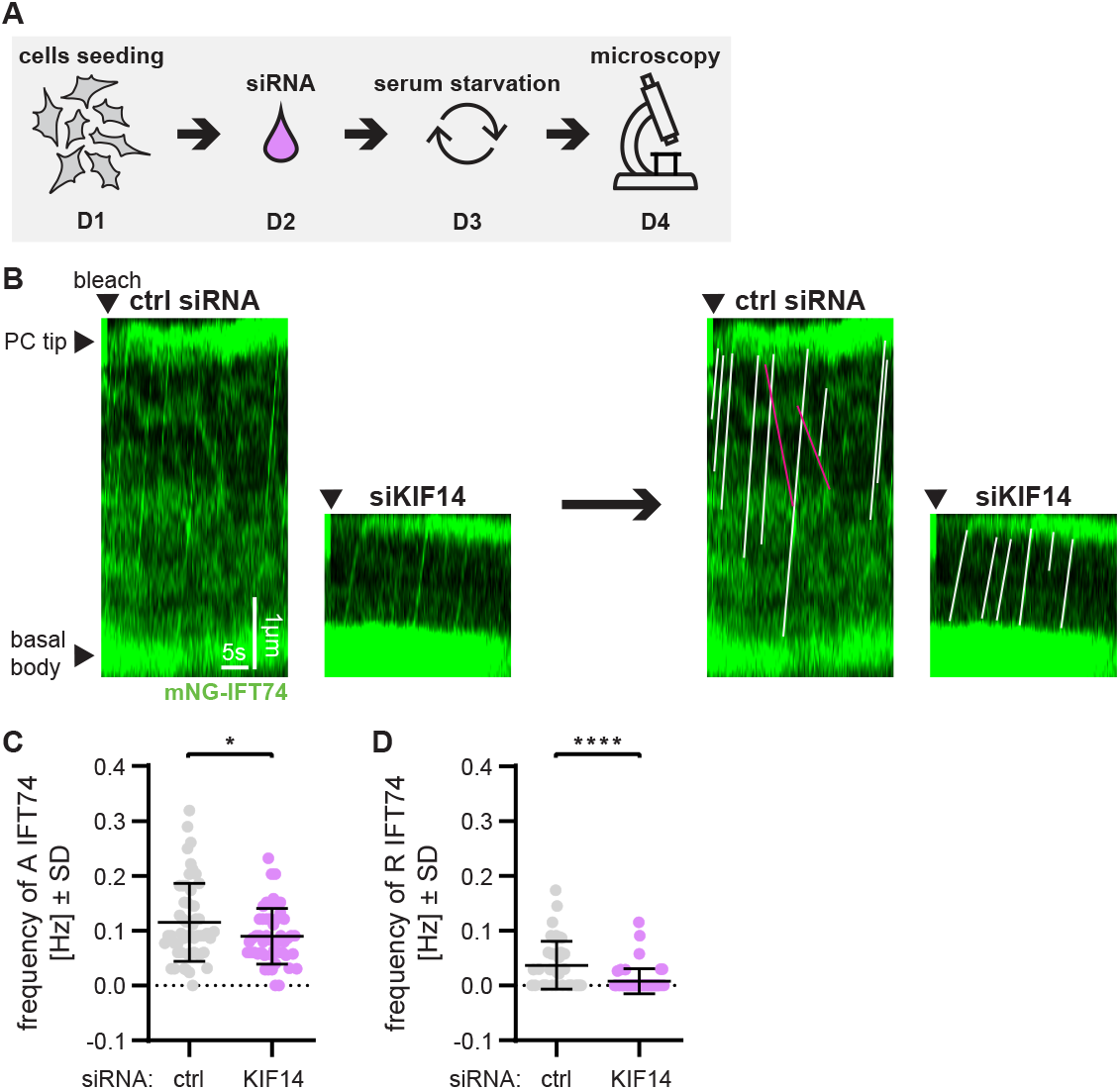
**(A)** Outline of the experiment from day(D)1 till day(D)4 to examine the effects of KIF14 depletion in hTERT RPE-1 cells. **(B)** Visualization of mNG-IFT74 transport events (anterograde movement indicated with white lines, retrograde with magenta) after KIF14 depletion using kymograph analyses; scale bar = 1 µm and 5 s, respectively. **(C-D)** Quantification of mNG-IFT74 transport events shows decreased frequency of anterograde (C) and retrograde (D) IFT74 trains after KIF14 depletion (unpaired t-test, *P < 0.05, ****P < 0.0001); n = 4, N = 47-51 (depicted datapoints represent values from individual cilia).

Next, we used mNeonGreen(mNG)-IFT74 hTERT-RPE-1 reporter cell line to examine the dynamics of IFT following KIF14 loss. Consistently with our previous data (17), KIF14 depletion in mNG-IFT74 hTERT-RPE-1 led to a prominent reduction of primary cilia length (Suppl. Fig. 1B). Using confocal live cell imaging, we examined the trafficking of mNG-IFT74 within cilia. First, we detected bi-directional movement of mNG-IFT74 particles (trains), corresponding to the anterograde and retrograde transport, respectively (Suppl. video 1 and 2). Using a kymograph analysis in ImageJ we visualized the IFT events and in turn tracked IFT trajectories clearly distinguishable from the background (Fig. 1B). Next, by analyzing the number of trajectories per time and their angle (30) we determined the corresponding velocities and frequencies of the IFT trains (see Material and methods for details). We found that IFT particles moved in the anterograde direction more frequently (0.12 Hz, Fig. 1C) and with a higher velocity (0.73 µm/s, Suppl. Fig. 1C) than in the retrograde direction (0.04 Hz, Fig. 1D and 0.40 µm/s, Suppl. Fig. 1D), in line with earlier observations (18,31,32). Having confirmed the observed trafficking parameters of mNG-IFT74 were within the range of values previously reported for vertebrate primary cilia, we subsequently examined the effects of KIF14 depletion. Intriguingly, while the velocity of mNG-IFT74 movement was comparable between control and KIF14-depleted cells, both in case of the anterograde (0.68 µm/s, Suppl. Fig. 1C) and the retrograde direction (0.49 µm/s, Suppl. Fig. 1D), the transport frequencies were significantly reduced in KIF14 depleted cells (0.09 Hz in the anterograde direction, 0.01 Hz in the retrograde direction, Fig. 1C-D). Together, our results suggested that KIF14 is required for correct IFT dynamics in the primary cilia of human cells.

### KIF14 controls intraciliary transport independently of AURA

Our earlier work revealed that KIF14 regulates primary cilia in both Aurora A kinase (AURA)-dependent and -independent manner (17). To address whether the effects of KIF14 on IFT transport require AURA activity, we analyzed mNG-IFT74 dynamics following KIF14 depletion and AURA inhibition (with 100nM TCS7010 for 24h), see Fig. 2A. Using kymograph analyses we visualized the IFT events (Fig. 2B). As expected, AURA inhibition increased primary cilia length in both control and KIF14-depleted mNG-IFT74 hTERT-RPE-1 cells (Suppl. Fig. 2A). Concomitantly, inhibition of AURA activity had a moderately positive effect on mNG-IFT74 transport frequency in control cells (Fig. 2C, Suppl. Fig. 2B). Importantly, although AURA inhibition restored cilia length in KIF14-depleted cells (Suppl. Fig. 2A), it did not rescue the defect in mNG-IFT74 transport frequency under the same conditions (Fig. 2C). In addition, we performed FRAP (fluorescence recovery after photobleaching) measurement in the ciliary tip of mNG-IFT74 hTERT-RPE-1. In line with the reduced rate of IFT, we detected a moderate decrease in signal recovery of mNG-IFT74 in KIF14-depleted cells (Fig. 2D-E). Interestingly, while KIF14 siRNA led to reduced signal recovery of mNG-IFT74 over control cells, inhibition of AURA activity had no visible effect on mNG-IFT74 recovery in KIF14-depleted cells. Moreover, forskolin treatment (100 μM, 24 hours), promoted cilia length in both control and KIF14-depleted mNG-IFT74 hTERT-RPE-1 cells (Suppl. Fig. 2C), in line with its reported positive effect on IFT velocity (33). Primary cilia length depends on several factors, including IFT velocity and frequency, the size/composition of IFT trains, the rate of IFT particle assembly and cilia entry, the interaction of IFT trains with their cargo, etc. (8,34). Consistent with the proposed models of cilia length regulation (34,35), our kymograph analysis of mNG-IFT74 transport indicates that cilia length and IFT frequency can be decoupled in hTERT-RPE-1 cells. We reason that the increase in cilia length of AURA-inhibited cells, without a concomitant effect on IFT frequency in KIF14-depleted cilia, could stem from an effect on train size and cargo loading (34–36), and/or altered turnover of axonemal tubulins (37,38) in AURA inhibited cells. In summary, our results demonstrated that some aspects of KIF14 regulation of IFT dynamics are not mediated by the effects of KIF14 on AURA, prompting us to investigate the responsible mechanism.

**Figure 2.**
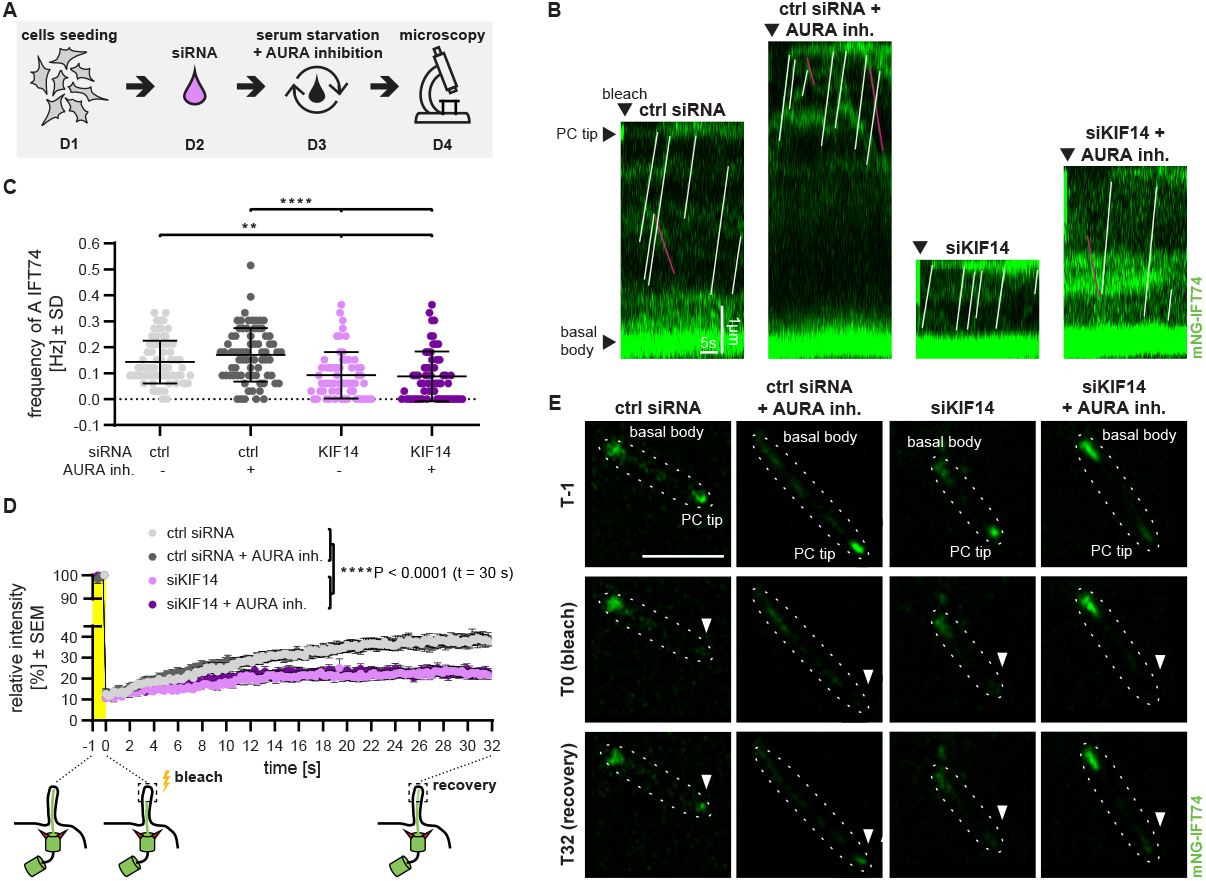
**(A)** Outline of the experiment from day(D)1 till day(D)4 to examine the effects of KIF14 depletion/AURA inhibition in hTERT RPE-1 cells. **(B)** Visualization of mNG-IFT74 transport events (anterograde movement indicated with white lines, retrograde with magenta) after KIF14 depletion and/or AURA inhibition using kymograph analyses; scale bar = 1 µm and 5 s, respectively. Quantification of mNG-IFT74 anterograde transport frequency is shown in **(C)** (one-way ANOVA, **P < 0.01, ****P < 0.0001); n=3, N = 60-80 (depicted datapoints represent values from individual cilia). **(D)** Measurement of recovery of mNG-IFT74 in the cilia tip using FRAP (one-way ANOVA, ****P < 0.0001); n = 3, N = 71-87. **(E)** Representative images from the FRAP experiment of mNG-IFT74. The primary cilium (PC) and surrounding area are outlined using a dashed line; scale bar = 5 µm.

### KIF14 displays IFT motor-like motility inside primary cilia

Given that our earlier work suggested that KIF14 depletion affects the localization of IFT57 at the cilia base (17), we tested whether KIF14 may control IFT dynamics via its effects on IFT protein basal body recruitment. However, we found no major alterations in mNG-IFT74 turnover at the cilia base using FRAP (Suppl Fig. 3A).

**Figure 3.**
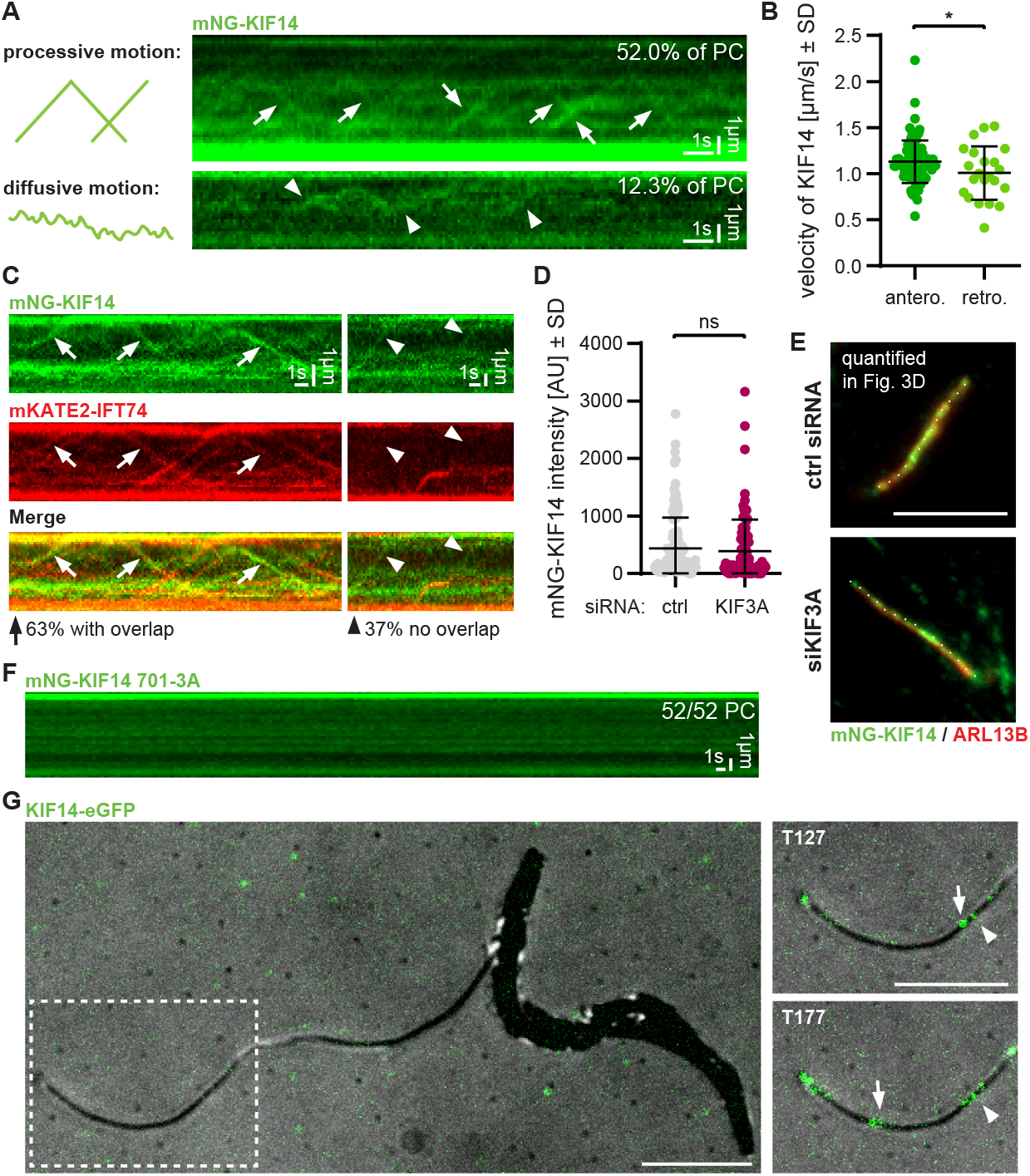
**(A)** Visualization of mNG-KIF14 processive (arrows) and diffusive (arrowheads) movements in primary cilia (PC) of hTERT-RPE-1 cells using kymograph analysis of live-cell TIRF microscopy experiments. The incidence of each type of motility (% of PCs) is shown; n = 1, N = 73, scale bar = 1 µm and 5 s, respectively. **(B)** Graph depicting measured velocities of mNG-KIF14 anterograde (antero.) and retrograde (retro.) processive movement (unpaired t-test, *P < 0); n = 1, N = 87-22, (depicted datapoints represent values from individual tracks). **(C)** Visualization of mNG-KIF14 and mKATE2-IFT74 movement in the cilium of hTERT-RPE-1 cells using kymograph analysis. 63% of processive mNG-KIF14 trajectories overlapped with those of mKATE2-IFT74 (arrows), while 37% of mNG-KIF14 processive trajectories lacked corresponding mKATE2-IFT74 signal (arrowheads); scale bar = 1 µm and 5 s, respectively, n = 1, N = 27. **(D)** Analysis of the mNG-KIF14 intensity in a primary cilium of hTERT RPE-1 cells after KIF3A depletion; n = 2, N = 90-125, (depicted data points represent values of individual cilia analyzed). **(E)** Localization of mNG-KIF14 in a primary cilium of hTERT RPE-1 cells after KIF3A depletion. ARL13B staining was used to detect primary cilia. The dashed lines illustrate the primary cilia, which were analyzed in D; scale bar = 5 µm. **(F)** Visualization of mNG-KIF14 701-3A (lack of) movement in primary cilia using kymograph analysis; n = 1, N = 52, scale bar = 1 µm and 5 s, respectively. **(G)** TIRF microscopy image of a detergent-extracted cytoskeleton of a Trypanosoma brucei cell with a partially detached flagellar cytoskeleton. TIRF microscopy images illustrating diffusive (arrowheads) and processive (arrows) motion of mNG-KIF14 molecules along Trypanosoma brucei axonemal microtubules in vitro in time (T-s); scale bar = 5 µm.

To investigate the mobility of KIF14 inside primary cilia, we stably expressed mNG-KIF14 in hTERT-RPE-1 cells and following serum starvation analyzed its behavior using live-cell Total Internal Reflection Fluorescence (TIRF) microscopy (39,40). We detected mNG-KIF14 readily localizing inside primary cilia (Suppl. Fig. 3B). Interestingly, we observed a bi-directional movement of mNG-KIF14 particles in 64% of ciliated cells. Specifically, kymograph analysis revealed that mNG-KIF14 moved processively in 52% of primary cilia or using only the diffusive type of movement in about 12% of cilia (Fig. 3A). We reason that mNG-KIF14 was predominantly present in (auto)inhibited state in those cilia showing only the diffusive movement, in line with similar behavior of other kinesins (41–45). We subsequently calculated the velocity of processive trajectories of mNG-KIF14. We found that mNG-KIF14 molecules moved in the anterograde and retrograde directions with an average velocity of 1.13 μm/s and 1.00 μm/s, respectively (Fig. 3B). This movement was significantly faster than the motion observed with GFP-KIF14 on purified microtubules *in vitro* (22), and also slightly faster than the velocity of mNG-IFT74 particles we determined in our earlier experiment using confocal microscopy (Suppl. Fig. 1C-D). Importantly, when re-measuring the anterograde and retrograde mNG-IFT74 velocities using the live-cell TIRF instrumentation (Suppl. Fig. 3C), we found them almost identical (1.07 μm/s and 0.98 μm/s, respectively) to that of KIF14. Next, we determined the frequency of mNG-KIF14 processive runs. As shown in Suppl. Fig. 3D, the obtained frequency of 0.02 Hz for anterograde and 0.004 Hz for retrograde transport was lower than the corresponding values of mNG-IFT74 (0.16 Hz in the anterograde direction and 0.07 Hz in the retrograde direction) determined using live-cell TIRF (Suppl. Fig 3E). These data raised an intriguing possibility that KIF14 and IFT74 may travel in close association in primary cilia, but also implied that IFT74 frequently moves from the base to the cilia tip and back without any associated KIF14 molecules.

Having found that KIF14 affects some aspects of IFT and moves in IFT-like fashion, we tested if IFT particles and KIF14 molecules move separately or together inside primary cilia. To this end, we generated hTERT-RPE-1 reporter line stably co-expressing either mNG-KIF14 or mNG-KIF14 738ΔP (please note this mutant is examined in detail in Fig. 4), and mKATE2-IFT74. Following serum starvation, we tracked the movement in primary cilia using live-cell TIRF microscopy. Interestingly, we observed that a majority of processive trajectories (63% for mNG-KIF14, 84% for mNG-KIF14 738ΔP) overlapped with those of mKATE2-IFT74 (Fig. 3C and Suppl. Fig. 3F). This aligns with our earlier observation that KIF14 and IFT74 move through primary cilia at similar velocities. Further, a fraction (37% for mNG-KIF14, 16% for mNG-KIF14 738ΔP) of KIF14 processive trajectories lacked the corresponding signal of mKATE2-IFT74 (Fig. 3C and Suppl. Fig. 3F).

**Figure 4.**
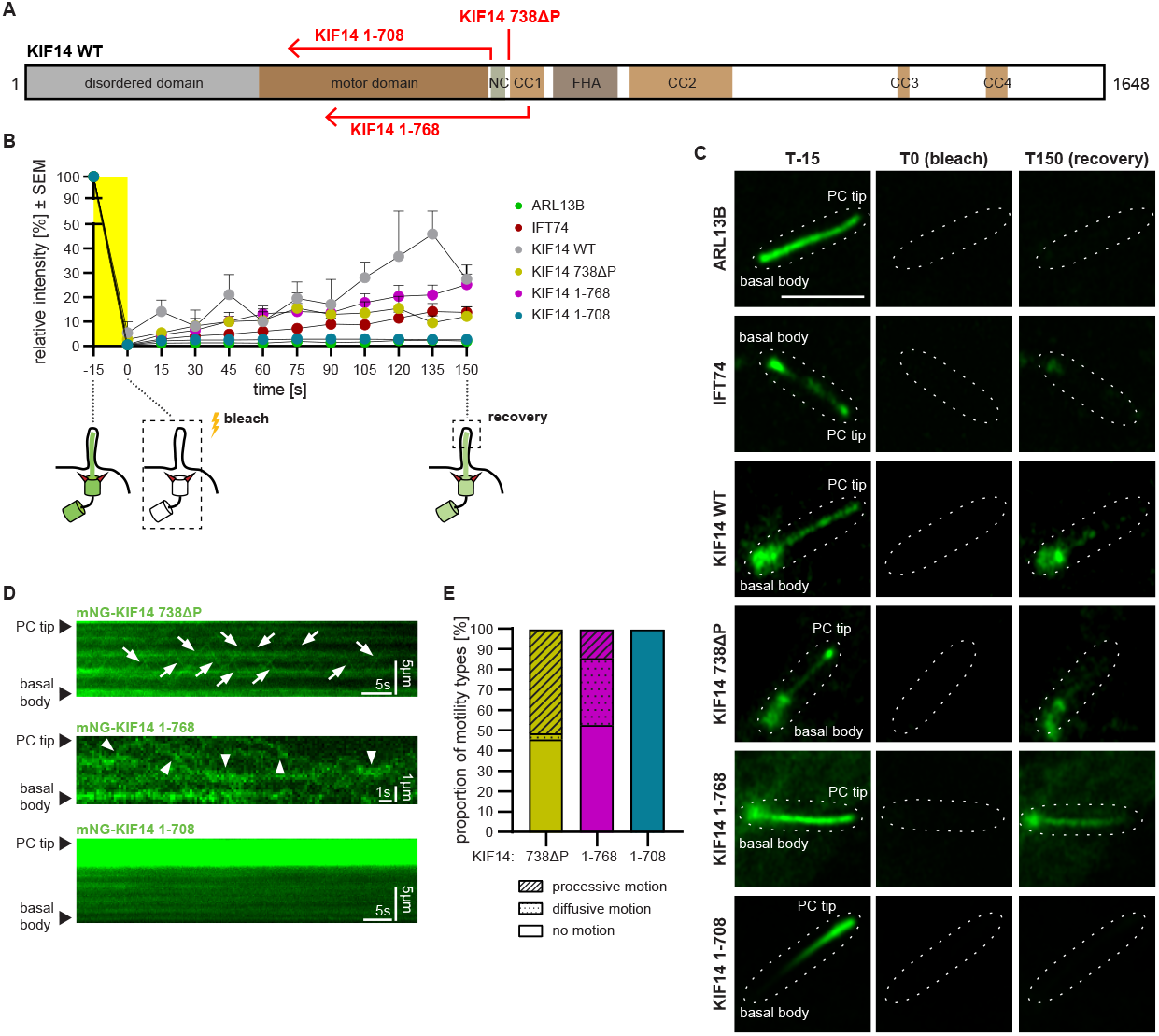
**(A)** Schematic representation of KIF14 functional domains in WT and the generated mutants (1-708, 1-768, 738ΔP). **(B)** FRAP analysis of mNG-tagged KIF14 (WT and mutants) following whole cilium bleach (see the scheme below the graph) in hTERT-RPE-1 cells. IFT74 and ARL13B were used as positive and negative controls, respectively; n = 1-3, N = 3-48. The corresponding representative images are shown in **(C)**. The primary cilium (PC) with surrounding area is outlined using a dashed line; scale bar = 3 µm. **(D)** Live-cell TIRF microscopy analysis of mNG-tagged KIF14 mutants in hTERT-RPE-1 cells, visualized using kymographs. Arrowheads point to diffusive movement, and arrows indicate processive movement; scale bar = 5 µm and 5 s, respectively. The incidence of each type of motility for the examined KIF14 mutants is shown in **(E)**; n = 2, N = 87-119.

Our observations suggest two possible explanations. First, KIF14 could be transported along with IFT proteins by another motor kinesin (most likely kinesin-2). The observed mKATE2-IFT74-negative kymograph trajectories of KIF14 movement may reflect its transport alongside endogenous (unlabeled) IFT trains or instances where the mKATE2-IFT74 signal photobleached before recording began. Indeed, a lack of motor domain in homodimeric KIF17 or motor activity in KIF7, immotile kinesin-4 regulating ciliary signaling, still allows their intraciliary movement via the action of other motors (46,47). Alternatively, KIF14 could rely on its motor domain to drive its movement inside the primary cilia and in this way potentially participate in the transport of a subset of IFT molecules or other cargoes. While a possible contribution of KIF13B to IFT is unclear, WT KIF13B (but not its motor mutant), is able to move processively in vertebrate cilia at velocities similar to IFT (18). To address whether KIF14 relies on its motor activity or the activity of other kinesins in cilia, we first examined ciliary localization of mNG-KIF14 in cells depleted of a subunit of kinesin-II, KIF3A. Importantly, we observed that KIF3A depletion decreased the percentage of ciliated cells (Suppl. Fig. 3G) and reduced IFT74 levels at the ciliary base (Suppl. Fig. 3H, J) and the axoneme (Suppl. Fig. 3I, J). However, it did not alter ciliary levels of mNG-KIF14 (Fig. 3D, E). This suggests that KIF14 ciliary localization does not require kinesin-II.

Next, we mutated residues 701-703 (I701A/V702A/N703A) of the KIF14 neck linker, adjacent to the globular core of the motor domain (48), corresponding to the motility-defective mutation in kinesin-1 (49), to generate a motor-defective version of KIF14 701-3A (Suppl. Fig. 3K). Importantly, we found that while mNG-KIF14 701-3A could localize to the primary cilia (Suppl. Fig. 3L), it did not show any motility in any of 52 primary cilia we analyzed, (Fig. 3F).

Purified KIF14 had been demonstrated to move in a directed manner along microtubules *in vitro* (22). To study whether it is also capable of moving on axonemal microtubules, we imaged the behavior of purified KIF14 on the flagellar cytoskeletons prepared by lysing cells of *Trypanosoma brucei*. Interestingly, we were able to detect both processive and diffusive motion of purified KIF14-eGFP along the axonemal microtubules *in vitro*, using single-molecule TIRF microscopy (Fig. 3G, Suppl. Fig. 3M, supplementary video 3). The subsequent kymograph analysis revealed the average velocity of KIF14-eGFP processive motion as 0.09 μm/s (Suppl. Fig. 3N). We noted that while the observed average velocity was in line with values reported for motility of purified KIF14 along stabilized microtubules *in vitro* (22), it was slower than the movement of KIF14 or IFT74 seen in primary cilia of hTERT-RPE-1, and significantly slower than the reported velocity (∼ 2.5 µm/s) of anterograde IFT in intact flagella of *Trypanosoma brucei* (50). The observed differences may arise from the specificities/customizations of these individual systems, including different states of modification of tubulin lattices (51,52). In addition, the effects of cargo binding and/or cooperation with other motors (53,54) may account for different velocities of KIF14 in vitro and in cells.

In summary, these results allowed us to draw several conclusions: i) KIF14 does not play a major role in the turnover of IFT74 complexes at the cilia base. Our data on IFT74 dynamics at the basal body align with the diffusion-to-capture model, where the majority of IFT74 and other IFT proteins are recruited from the cytoplasm to the basal body to assemble into IFT trains, and within 5-20 seconds enter the cilium (55–57). ii) KIF14 has a role in intraciliary transport, by being able to perform both processive and diffusive types of movement in primary cilia of hTERT-RPE-1 and walk on purified axonemes of *Trypanosoma brucei*. Importantly, KIF14 cilia localization does not rely on KIF3A, a subunit of kinesin-II. This makes KIF14 the fourth kinesin (after KIF3A/3B complex, KIF17, and KIF13B (16)) capable of processive movement in vertebrate primary cilia. iii) KIF14 trajectories often overlap with trajectories of IFT particles, implying their close association. KIF14 could act as cargo of IFT complex or could contribute to IFT through its motor activity. The former scenario offers a reasonable explanation for the observed retrograde movement of KIF14 molecules inside cilia, likely driven by dynein-2-dependent retrograde IFT. However, in contrast to the motor-defective KIF17 mutant (46) and rigor (a mutant with strong binding to microtubules) and microtubule-binding-defective KIF7 mutants (47), the KIF14 701-3A motor mutant shows a prominent motility defect in cilia. Thus, it seems highly plausible the observed anterograde movement of KIF14 is indeed mediated by the activity of its motor domain, rather than simply being transported in a “cargo-like fashion” by anterograde IFT complexes like some of the other non-canonical ciliary kinesins. iv) KIF14 seems to control IFT dynamics by regulating the transport of a subset of IFT particles, making it the first reported non-canonical kinesin modulating IFT in the primary cilia of vertebrate cells. In this model, such activity could arise from its role in IFT train assembly and/or its direct participation in propelling some IFT particles or other cargoes via the actions of its motor domain. As for the latter, the role of KIF14 in driving the anterograde IFT transport would be supplementary to the major role of kinesin-2 and possibly being cell type-specific. This is in line with the moderate severity of KIF14-depletion phenotypes for IFT (compared to the dire consequences of kinesin-2 ablation (10–14,16)) as well as the model of multiple molecules of kinesin motor binding simultaneously to an IFT train (53,54). We note that ultimately linking KIF14 to IFT regulation would require resolving its position within native IFT particles using a structural biology approach, a feat not yet accomplished even for the canonical ciliary kinesin, kinesin-II (6). v) As KIF14 is a plus-end-directed motor, its anterograde motility should reflect the activity of its motor domain, while its retrograde transport in cilia is likely mediated by other means. For instance, kinesin-2 molecules can after dissociation from IFT complexes at the cilia tip either travel back to the cilia base by diffusion in *Chlamydomonas* (58) or are moved as cargo by the dynein-powered retrograde IFT complexes in cilia of *C. elegans* (54). What is the situation in vertebrate cilia is far less clear. Our data on KIF14 701-3A show that the neck linker mutation impairs both anterograde and retrograde motility of KIF14. A similar defect was reported for a motor domain mutant of KLP-6 (ortholog of KIF13B) in *C. elegans* (19). This may reflect the rigor state of the corresponding motor mutants (strong association with microtubules due to the inability to hydrolyze ATP (48)), but we cannot formally exclude that the motor activity of KIF14 affects its retrograde transport and/or its interaction with IFT complexes, in line with the effects of cargo-kinesin interactions on motor processivity (45,59). These observations warrant a more detailed investigation of the mechanism of KIF14 in primary cilia and its interactions with IFT complexes in the future.

### C-term regions of KIF14 are critical for its motility in primary cilia

KIF14 consists of a disordered N-terminal region, motor domain, and C-terminal part containing coiled-coil domains and unstructured regions (Fig. 4A). While the ability of kinesins to move along the microtubules is primarily determined by the activity of the motor domain, the nature of the motion is also affected by both intermolecular and intramolecular interactions of the C-terminal region (44). These interactions may regulate the processivity of the movement, dimerization of kinesin molecules, interactions with cargo, etc. (22,42,44). To determine the role of the C-terminal part of KIF14 in its movement along ciliary microtubules, we generated several KIF14 mutants. Constructs 1-708 and 1-768 lacked the whole C-term part, including the FHA domain, coiled-coil domains, and unstructured regions (Fig. 4A). In addition, construct 1-708 lacked a region implicated in the dimerization of KIF14 *in vitro* (22). Earlier work proposed that parallel dimerization of NC and CC1 regions inhibits processivity of several members of the kinesin-3 family (42,60), which can be relieved by a deletion of the first Proline of CC1 domain (42). Using AlphaFold structure prediction we pinpointed the NC and CC1 domains into 711-731 and 738-789 regions, respectively (Fig. 4A, Suppl. Fig. 4A). We deleted Pro 738 to generate KIF14 738ΔP construct to examine the role of this proposed mechanism in the mobility of KIF14 inside primary cilia.

Following the generation of reporter hTERT-RPE-1 lines for each mNG-tagged KIF14 variant, we observed all constructs readily localizing in primary cilia (Suppl. Fig. 4B-D), suggesting that the introduced changes did not dramatically affect ciliary targeting of individual KIF14 mutants. Noteworthy, we observed a prominent accumulation of mNG-KIF14 1-708 in the distal part of the ciliary axoneme (Suppl. Fig. 4D). Next, we analyzed the dynamics of KIF14 mutants inside primary cilia using FRAP (Fig. 4B-C). We used GFP-ARL13B as an example of a ciliary component with slow turnover (61) and mNG-IFT74 as an example of a highly mobile ciliary component (62). Intriguingly, while mNG-KIF14 1-708 showed virtually no recovery after whole cilium bleach, mNG-tagged KIF14 wild type (WT), 1-768, and 738ΔP fusion proteins demonstrated prominent recovery.

To corroborate this finding, we analyzed the movement of KIF14 C-term mutants using live-cell TIRF microscopy. We found that both mNG-KIF14 1-768 and mNG-KIF14 738ΔP were able to move in 47% and 54% of ciliated cells (Fig. 4D-E), respectively, in line with the behavior of full-length KIF14 we analyzed earlier (Fig. 3A). KIF14 1-768 demonstrated diffusive movement (33% of cilia) or processive movement (14% of cilia). KIF14 738ΔP showed more often a processive type of motion (51% of cilia), and it showed only diffusive motion in 3% of primary cilia. However, we found no major difference in the frequency of processive runs between mNG-KIF14 and mNG-KIF14 738ΔP (Suppl. Fig. 4E-F), implying the NC-CC1 dimerization does not play a major role in the regulation of KIF14 motility in cilia. In agreement with no FRAP recovery of mNG-KIF14 1-708, we did not detect any of its processive or diffusive motion inside primary cilia (Fig. 4D-E). However, we observed the diffusive motion of mNG-KIF14 1-708 along the cytosolic microtubules (Suppl. Fig. 4G-H). In addition, using single-molecule TIRF microscopy, we found mNG-KIF14 1-708 from cell extracts was capable of slow processive runs on GMPCPP-stabilized microtubules *in vitro* (Suppl. Fig. 4I). Here, we also detected accumulation of mNG-KIF14 1-708 at the ends of GMPCPP-stabilized microtubules, in agreement with our observations from cells (Suppl. Fig 4D), but in contrast to mNG-KIF14 WT used as a control (Suppl. Fig. 4J-K). In sum, the data above demonstrated that C-terminal regions of KIF14 are critical for the protein motility and dynamics in primary cilia. One possible explanation of the defects seen with KIF14 1-708 is impaired dimerizing, which is known to significantly impact kinesin processivity (44). Indeed, KIF14 1-708 fails to efficiently dimerize *in vitro* (22), which seems to render it immobile (“rigor-like”) in primary cilia, but not on cytosolic microtubules. In fact, our data showing different behavior of mNG-KIF14 1-708 in primary cilia and on cytosolic microtubules provide a plausible example of how the motor capabilities could be tailored by interactions with specific types of lattice and its tubulin modifications (51,52).

### Motility of KIF14 in primary cilia requires CC1 rather than NC domain

Having found that the region between 708 and 768, encompassing the NC and CC1 domains, has a profound impact on the motility and dynamics of KIF14 in primary cilia, we generated KIF14 1-768 ΔNC (lacking 709-735 region) and KIF14 1-768 ΔCC1 (lacking 736-768 region) mutants (Fig. 5A). Interestingly, we observed that mNG-KIF14 1-768 ΔCC1 mimicked the behavior of mNG-KIF14 1-708. In contrast to mNG-KIF14 1-768 ΔNC and mNG-KIF14 1-768, it showed a prominent accumulation in the distal part of the cilium in hTERT-RPE-1 cells (Suppl. Fig. 5A-B). Moreover, it failed to show any reasonable recovery following the whole cilium bleach in FRAP experiments (Fig. 5B-C). Thus, this data suggested that CC1, but not the NC domain, is critical for the dynamics and motility of KIF14 in primary cilia.

**Figure 5.**
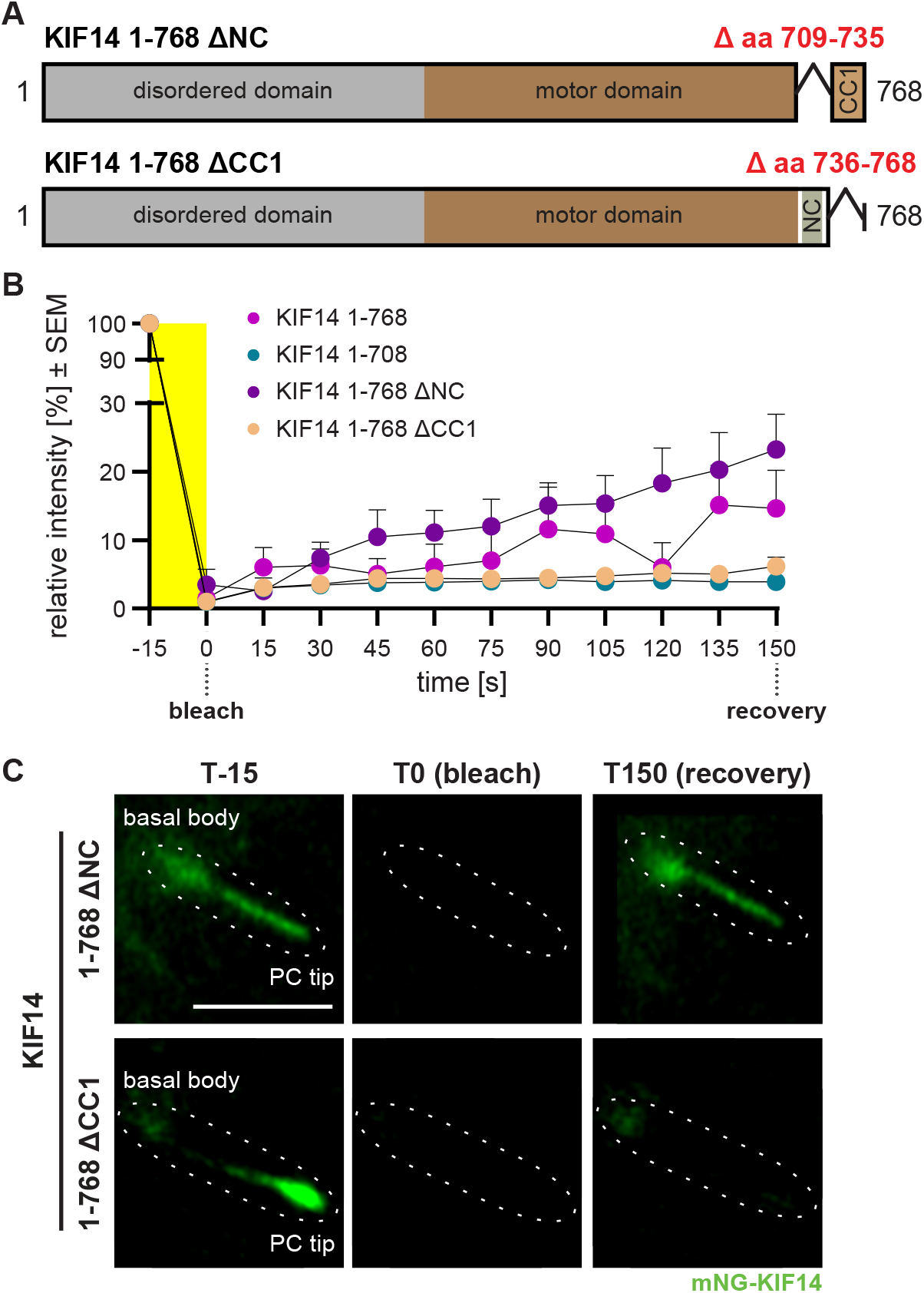
**(A)** Schematic representation of KIF14 1-768 ΔNC and ΔCC1 mutants. **(B)** FRAP analysis of mNG-tagged KIF14 mutants (1-708, 1-768, 1-768 ΔNC, and ΔCC1) following whole cilium bleach in hTERT-RPE-1 cells; n =3, N = 7-19. The corresponding representative images are shown in **(C)**. The primary cilium (PC) with the surrounding area is outlined using a dashed line; scale bar = 3 µm.

Considering the reported roles for the NC-CC1 interface in other kinesin-3 family members, this was an unexpected observation. In KIF1A and KIF16B, deletion of the CC1 domain notably improves the processivity of a given motor, while the NC domain is crucial for dimerization (42), and in turn processivity. Moreover, similar effects were reported for KIF13B (60). Thus, the requirement for CC1 for protein motility makes KIF14 unique among other kinesin-3 members. Intriguingly, examination of AlphaFold-predicted NC-CC1 regions revealed notable differences between CC1 of KIF14 and that of the other kinesin-3 members. While the regions corresponding to CC1 were either short, fragmented, and/or of low confidence of coiled-coil, the predicted structure of CC1 of KIF14 showed a coiled-coil region with the highest confidence and length (Suppl. Fig. 5C). As kinesin dimerization is intimately linked to its processivity and dynamics, we speculate that the mechanism of KIF14 dimerization is different from other family members, by relying on CC1 rather than NC domain. This model warrants in-depth examination of KIF14 dimerization in cilia by structural biology approaches in the future.

### C-term KIF14 mutants impair IFT by causing traffic jam-like accumulations in primary cilia

Our data suggested that lack of CC1 domain impairs the motility of KIF14 inside cilia, leading to enrichment of the mutated protein in the distal part of the cilium. As we observed that KIF14 modulates IFT, we next examined the state of IFT in such cilia. First, we noted that enrichment of mNG-KIF14 1-708 in the distal parts of cilia led to a concomitant accumulation of IFT88 and ARL13B (Suppl. Fig. 6A-B), and in turn enlarged cilia tips (particularly visible using ARL13B staining; Suppl. Fig. 6A) reminiscent of bulged cilia due to impaired transport from the cilia tip to the base (9). To assess a possible impairment of ciliary trafficking, we examined the movement of mKATE2-IFT74 complexes inside those cilia by live-cell TIRF microscopy. Interestingly, we noted that mNG-KIF14 1-708 had a profound negative effect on the frequency of both anterograde and retrograde IFT when compared to mNG-KIF14 WT (Fig. 6A-B). Specifically, the anterograde IFT74 frequency dropped from 0.21 Hz to 0.07 Hz, and the retrograde frequency decreased from 0.08 Hz to 0.01 Hz. Similarly, we detected a small but consistent drop in anterograde (from 1.15 µm/s to 1.08 µm/s) and retrograde (from 1.04 µm/s to 0.91 µm/s) IFT velocity in cilia of mNG-KIF14 1-708 expressing cells (Fig. 6C-D). In addition, we observed an increased incidence of trafficking artifacts such as stopping trains (14% in WT versus 36% in KIF14 1-708, Fig. 6E). We suspect the rather high level of transport errors we detected in our control (KIF14 WT) may be due to overexpression of KIF14 and IFT proteins above physiological levels, and/or reflect the variable length of axonemal microtubules typical for primary cilia of vertebrate cells (63). In any case, our data strongly suggested that KIF14 1-708 mutant causes defects in IFT in primary cilia.

**Figure 6.**
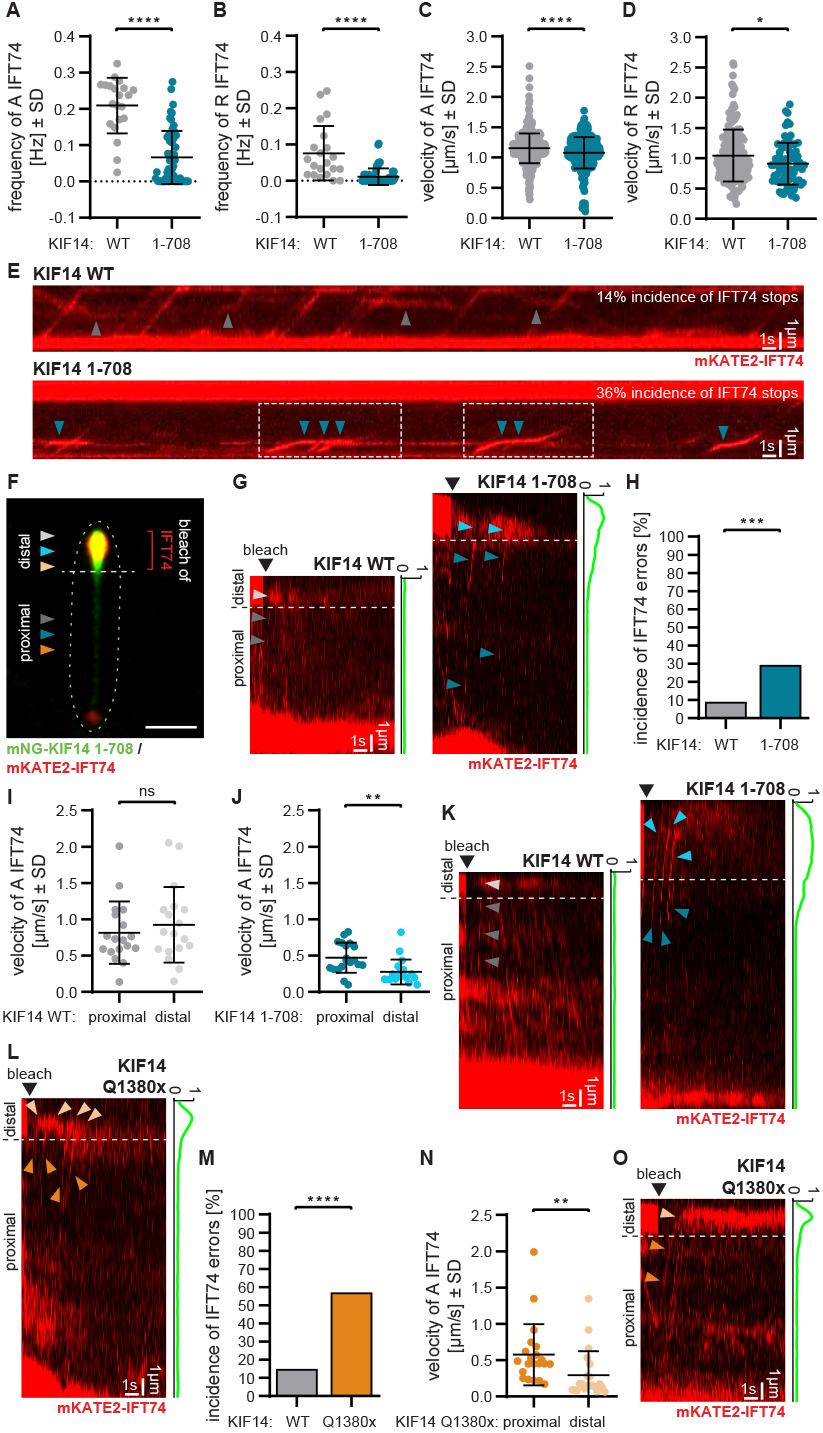
**(A-D)** Live-cell TIRF microscopy analysis of mKATE2-IFT74 transport events (movement of anterograde or retrograde trains) in the presence of mNG-KIF14 or mNG-KIF14 1-708 in hTERT-RPE-1 cells. Graphs demonstrate a reduction in the frequency of mNG-IFT74 anterograde trains **(A)**, retrograde trains **(B)**, and their anterograde **(C)** and retrograde **(D)** velocities in mNG-KIF14 1-708 containing cilia (unpaired t-test, *P < 0.05, ****P < 0.0001); n = 1, N = 21-56 for A and B (depicted datapoints represent values from individual cilia), N = 76-525 for C and D (depicted datapoints represent values from individual tracks). The corresponding kymograph analysis is shown in **(E)**, including the percentage of incidence of mKATE2-IFT74 stops (gray arrowheads for KIF14 WT and blue arrowheads for KIF14 1-708); n = 1, N = 411-634, scale bar = 1 µm and 1 s, respectively. Dashed rectangles highlight a situation we termed “trains in tandem” (IFT trains sequentially stopping at the same location, to eventually resume transport as one merged train) **(F)** Schematic of the experiment for tracking and measuring mKATE2-IFT74 velocities in proximal (dark gray arrowhead for KIF14 WT and dark blue arrowhead for KIF14 1-708) and distal (light gray arrowhead for KIF14 WT and light blue arrowhead for KIF14 1-708) parts of primary cilia after bleaching of mKATE2-IFT74 in the distal part of the cilium. The boundary between the proximal and distal parts was determined based on the accumulation of mKATE2-IFT74 (and mNG-KIF14/KIF14 1-708) and is outlined with a dashed line; scale bar = 2 µm, **(G)** Examples of mKATE2-IFT74 not reaching the tip in KIF14 WT (dark gray arrowheads for proximal part and light gray arrowheads for distal part of primary cilium) and 1-708 (dark blue arrowheads for proximal part and light blue arrowheads for distal part of primary cilium). The boundary between the proximal and distal parts is outlined with a dashed line; scale bar = 1 µm and 1 s, respectively. The plot profile next to the kymograph (in G and K) illustrates the distribution of mNG-KIF14 WT relative to mNG-KIF14 1-708 along the primary cilium. **(H)** Elevated incidence of mKATE2-IFT74 not reaching the tip in KIF14 1-708 (29%) compared to KIF14 WT (9%) (unpaired t-test, ***P < 0.001); n = 3, N = 11-17. Velocities of IFT74 anterograde trains in the proximal and distal parts of the cilium in cells expressing KIF14 WT **(I)** and KIF14 1-708 **(J)** (paired t-test, **P < 0.01); n = 3, N = 18-19 (depicted datapoints represent values from individual tracks). **(K)** Kymograph visualization showing mKATE-IFT74 slowing down in the distal part containing high density of KIF14 1-708 (dark blue arrowheads for proximal part and light blue arrowheads for distal part of primary cilium) tip compared to KIF14 WT (dark gray arrowheads for proximal part and light gray arrowheads for distal part of primary cilium; scale bar = 1 µm and 1 s, respectively. **(L)** An example of mKATE2-IFT74 not reaching the tip in KIF14 Q1380x (dark orange arrowheads for the proximal part and light orange arrowheads for the distal part of the primary cilium). The boundary between the proximal and distal parts is outlined with a dashed line; scale bar = 1 µm and 1 s, respectively. The plot profile next to the kymograph (in L and O) illustrates the relative distribution of mNG-KIF14 Q1380x along the primary cilium. **(M)** Elevated incidence of mKATE2-IFT74 errors in reaching the tip in KIF14 Q1380x (64%) compared to KIF14 WT (15%) (unpaired t-test, ****P < 0.0001); n = 3, N = 13-63. **(N)** Velocities of mKATE2-IFT74 anterograde trains in the proximal and distal parts of the cilium in cells expressing KIF14 Q1380x (paired t-test, **P < 0.01); n = 3, N = 8-21 (depicted datapoints represent values from individual tracks). **(O)** Kymograph visualization showing mKATE-IFT74 slowing down in the distal part of the KIF14 Q1380x containing cilium (dark orange arrowheads for the proximal part and light orange arrowheads for the distal part of the primary cilium); scale bar = 1 µm and 1 s, respectively.

The precise tracking of mKATE2-IFT74 trains along the entire axoneme was hindered by its accumulation with mNG-KIF14 1-708 in the distal part of the cilium. To overcome this limitation, we subsequently tracked mKATE2-IFT74 after specifically bleaching the accumulated pool of mKATE2-IFT74 in the distal parts of cilia (Fig. 6F). While mKATE2-IFT74 trains typically reached the cilium tip in mNG-KIF14 WT containing cilia, the incidence of errors of not reaching the tip (Fig. 6G) was markedly elevated in the presence of mNG-KIF14 1-708 (29% versus 9%, Fig. 6H). Remarkably, the mKATE2-IFT74 trains detected in the presence of mNG-KIF14 1-708 in the distal parts of the cilium displayed a prominent drop in their velocities (from 0.47 µm/s to 0.28 µm/s), compared to cilia containing mNG-KIF14 WT (0.82 µm/s and 0.92 µm/s, Fig. 6I-J). This abrupt change in the velocity of mKATE2-IFT74 particles was well noticeable in kymographs from movies obtained by both confocal (Fig. 6K) and live-cell TIRF imaging (Suppl. Fig. 6C) and coincided with the increase in mNG-KIF14 1-708 density. Even though kinesin-2 as the main anterograde IFT motor was proposed to be fairly resistant to self-crowding (64), we speculate that accumulation of KIF14 1-708 in the distal part of the cilia causes a highly crowded environment decreasing the IFT velocity by effectively creating a traffic jam (65). According to this model, the movement of IFT trains is hindered by the excess of KIF14 protein moieties, which are highly affinitive to axonemal microtubules and, in turn, represent a roadblock. Rigor mutants of kinesins were indeed demonstrated to act as efficient roadblocks *in vitro* (66). This aligns with reports that KIF14 lacking its C-terminal parts can associate with microtubules in a very stable “rigor-like” state *in vitro* (48). To our knowledge, this would be the first example of ciliary function being hampered by molecular crowding and subsequent traffic jams stemming from an accumulation of dysfunctional motors in cilia.

The observation of the “traffic jam”-like phenomenon and bulged cilia phenotype linked to the KIF14 C-term mutant led us to question its potential (patho)physiological relevance. Interestingly, *KIF14* mutations in patients can lead to C-terminally truncated proteins (29) which accumulate in the distal part of primary cilia (17). Given the similarity to the KIF14 1-708 localization along the cilium, we selected KIF14 Q1380x as an example of a disease-related mutation and asked if its presence also hampers ciliary trafficking, in a manner similar to KIF14 1-708. Indeed, we found that expression of mNG-KIF14 Q1380x led to an accumulation of IFT88 in distal parts of cilia (Suppl. Fig. 6D-E). Moreover, in agreement with the effects of KIF14 1-708, we observed a prominent increase in the IFT aberrant behavior (mKATE2-IFT74 particles were not able to reach the tip in 57% of analyzed tracks, Fig. 6L-M), as well as a notable slowdown of IFT of mKATE2-IFT74 particles (from 0.58 µm/s to 0.27 µm/s) moving in mNG-KIF14 Q1380x high-density regions in the distal part of cilia (Fig. 6N-O). In sum, our data demonstrated that the disease-related mutant KIF14 Q1380x significantly impairs ciliary trafficking, similar to the effects of KIF14 1-708. While the effects of examined KIF14 mutations may be facilitated by their exogenous levels in hTERT-RPE-1 cells, we found mNG-KIF14 Q1380x expressed at a level close to that of endogenous KIF14 (Suppl. Fig. 6F). Thus, the results suggest an attractive possibility that IFT defects due to traffic jams of dysfunctional motors may represent a mechanism underlying some of the KIF14-related pathologies. Given the differences in the C-term region between KIF14 Q1380x and KIF14 1-708, we cannot exclude that the underlying mechanisms making these mutants prone to accumulate in the distal part of cilia may differ. Future work should address the underlying biophysical aspects of this phenomenon as well as its *in vivo* relevance, together with the general role of traffic jams in defects of ciliary trafficking implicated in ciliopathies.

## Material and methods

### Cell culture, transfections, and small molecule treatment

hTERT RPE-1 cell line (gift from E. Nigg, Biozentrum, University of Basel, Basel, Switzerland) and derived hTERT RPE-1 reported cell lines were cultured in DMEM/F12 medium (Thermo-Fisher Scientific/Gibco) supplemented with 10% fetal bovine serum (FBS), 2 mM L-Glutamine, and 100 IU/mL Penicillin-Streptomycin (all from Biosera). Phoenix-Ampho packaging cell line (obtained from ATCC) was cultivated in high glucose glutamax DMEM medium (Thermo-Fisher Scientific/Gibco) supplemented with 10% fetal bovine serum (FBS), 2 mM L-Glutamine, and 100 IU/mL Penicillin-Streptomycin. All cell lines were maintained at 37 °C in a humidified atmosphere containing 5% CO_2_.

For siRNA experiments, hTERT RPE-1 cells were seeded to achieve approximately 80% confluency the next day. They were then transfected with 50 nM control siRNA or 50 nM KIF14 siRNA, or 10 nM KIF3A siRNA using 1.1 µl Lipofectamine RNAiMAX (Thermo-Fisher Scientific/Invitrogen) mixed in 50 ul OPTIMEM (Thermo-Fisher Scientific/Gibco). For KIF14 knockdowns, cells were transfected with a mixture of two siRNA oligos (50nM total; 25 nM each of oligo #1 and #2). The medium was replaced with a fresh one 24 hours after transfection. The siRNAs used are listed in Table 1.

To promote ciliogenesis, cells were starved in FBS-free DMEM/F12 medium supplemented with 2 mM L-Glutamine and 100 IU/mL Penicillin-Streptomycin for 24 hours. For live cell imaging, the medium was replaced with FBS-free FluoroBrite DMEM (Thermo-Fisher Scientific/Gibco) up to 24 hours prior to imaging.

For experiments involving AURA inhibition, hTERT RPE-1 cells were treated with 100 nM TCS7010 (Bio-Techne/Tocris Bioscience) AURA inhibitor 24 hours post-siRNA transfection, and the treatment was maintained for 24 hours.

In the experiment involving forskolin treatment, hTERT RPE-1 cells were treated with 100 μM forskolin (Sigma-Aldrich) 24 hours after siRNA transfection, and the treatment was maintained for 24 hours.

### DNA constructs and stable cell lines

Constructs encoding mutant forms of human KIF14 (738ΔP, 1-768, 1-708, 1-768 ΔNC, 1-768 ΔCC1, 701-3A, and Q1380x) were generated by site-directed mutagenesis of pENTR KIF14 entry vector, using specific forward and reverse primers (Table 1) and Quickchange II XL kit (Agilent), according to manufacturer’s instructions. The pENTR KIF14-FL vector was created by pENTR/D-TOPO Gateway cloning (Thermo-Fisher Scientific/Invitrogen) from a KIF14 human cDNA plasmid generously provided by F. Barr (University of Oxford, Oxford, UK; (24)). Expression vectors were then generated using LR Gateway cloning (Thermo-Fisher Scientific/Invitrogen) to transfer the individual inserts (WT and mutant KIF14, and IFT74) from pENTR/D-TOPO vector into pMSCV-N-mNG-IRES-PURO and pMSCV-N-mKATE2-IRES-PURO destination vectors, respectively.

hTERT RPE-1 mNG-KIF14 cell lines (WT, 738ΔP, 1-768, 1-708, 1-768 ΔNC, 1-768 ΔCC1, 701-3A, and Q1380x), hTERT RPE-1 mNG-IFT74 cell line, or hTERT RPE-1 mNG-KIF14 (WT, 1-708 or Q1380x) co-expressing mKATE2-IFT74 were generated by retroviral transduction. Retroviruses were produced using the Phoenix-Ampho cell line seeded in a 25 cm^2^ culture flask. The highest transfection efficiency was achieved by transfecting the cells the following day, once they reached 80% confluency. On that day, the medium was replaced with 3 ml of Phoenix transfection medium, and cells were transfected with 8 µg of pMSCV-mNG vector containing the gene of interest (GOI) in 300 µl OPTI-MEM using 20 µl of 2x polyethylenimine (PEI). After 3 hours, the medium was replaced with 3 ml of viral particle production medium. Viral particle-enriched medium was collected every 12 hours for a total of 36 hours. This medium was filtered using 0.45 µm low-binding syringe filters (Filtratech). The filtered viral particle-enriched medium was immediately used to transduce RPE-1 cells using the spinfection method (30). Target RPE-1 cells were seeded in a 6- or 12-well plate one day prior to transduction to achieve approximately 80% confluency. The medium was discarded, and 1-2 ml of the freshly prepared viral particle-containing medium, supplemented with 10 µg/ml polybrene (Merck/Sigma-Aldrich), was added to the RPE-1 cells. The parafilm-sealed plate was then spinfected at 1100 x g for 45 minutes at room temperature (RT). After the overnight transduction, cells were washed with PBS and a fresh pre-warmed RPE-1 complete medium was added. GFP-ARL13b reporter cell line was generated by integration of pgLAP1_Neo – ARL13B into hTERT-RPE-1 Flp-In™ T-REx™ cells (30).

Cells were subsequently sorted via FACS and expanded either as single-cell clones or as mixed populations. Validation of stable transgene expression was performed using IF microscopy or Western blotting.

### Immunofluorescence staining

Cells were cultured on coverslips for 2-4 days, depending on the design of a particular experiment. The cells were then fixed with cold methanol for 10 minutes at -20°C, washed with PBS, and blocked in a blocking buffer (1% BSA in PBS) for 10 minutes at RT. Primary and secondary antibodies (listed in Table 1) were applied in blocking buffer for 1 hour at RT, each time followed by 3 PBS washes. Nuclei were stained with Hoechst for 5 minutes at RT. Following a final PBS wash, the coverslips were mounted using ProLong™ Glass Antifade Mountant (Thermo-Fisher Scientific).

Microscopy analyses of fixed cells were performed on a Zeiss AxioImager.Z2 equipped with a Hamamatsu ORCA Flash 4.0 camera. Z-stack images were acquired using 100x/1.4 Plan-Apochromat oil immersion objective, controlled by Zen Blue software. Z-slices were projected into a single layer using maximal intensity Z-projection, and color composite images were created in ImageJ (67), version 1.54k). For representative images, deconvolution was performed using Zen Blue software with a strength setting of 9 to 10 and up to a maximum of 40 iterations.

Cilia length was measured in ImageJ by tracing the ARL13B fluorescent signal with a segmented line. Typically, 15-20 cilia were measured per condition and experiment.

Intensity signal measurements for mNG-KIF14 (Fig. 3D) and IFT74 (Suppl. Fig. 3H-I) were performed using ImageJ. Z-projected images were processed with consistent brightness and contrast settings. A segmented line was drawn along the primary cilium from base to tip (Suppl. Fig. 3J), and the mean intensity values (in arbitrary units) were recorded. IFT74 signal at the ciliary base was manually outlined with a circle of uniform size for each primary cilium (Suppl. Fig. 3J), and the mean intensity values (in arbitrary units) were recorded. About 85-125 cilia were analyzed per condition and experiment Area measurements for KIF14, IFT88, and ARL13B (Suppl. Fig. 6B and 6E) were performed using ImageJ. Z-projected images were processed with consistent brightness and contrast settings. The area (in square pixels) containing the visible signal of each protein at the ciliary tip was manually outlined, and the mean intensity values were recorded. About 20-30 cilia were analyzed per condition and experiment.

Signal profiles of mNG-KIF14 WT/1-708/Q1380x (Fig. 6G, 6K, 6L, 6O, and Suppl. Fig. 6C) were analyzed using ImageJ. A segmented line was drawn along the primary cilium from base to tip. The measured values (mean signal intensity in arbitrary units) for individual cilia were plotted as histograms (with X-axis showing the cilia length in μm, and Y axis showing the measured intensities). In Fig. 6G, 6K, 6L, 6O, and Suppl. Fig. 6C, the data were normalized relative to the signal of mNG-KIF14 1-708/Q1380x (with the highest value of mNG-KIF14 1-708/Q1380x set to 1).

### Time-lapse live cell imaging using confocal microscopy

Time-lapse live cell imaging analysis of mNG-IFT74-positive primary cilia (Fig. 1B-D, Suppl. Fig. 1B-D, Fig. 2B-C, Suppl. Fig. 2A-B, Fig. 6F-O) was done as described before (30). Briefly, cells were seeded on IBidi ibiTreat m-Slide 8-well chamber slide (IBidi), where indicated (Fig. 1B-D, Suppl. Fig. 1B-D, Fig. 2B-C, Suppl. Fig. 2A-B), transfected with siRNA and/or treated with TCS7010 AURA inhibitor, and starved using FBS-free FluoroBrite imaging DMEM.

Time-lapse imaging of mNG-IFT74 was conducted using an inverted Zeiss LSM880 laser scanning confocal microscope mounted on an Axio Observer.7 stand, equipped with an Airyscan detector and Monochromatic camera Axiocam MRm, CCD sensor, 1388 × 1040 pixels, pixel size 6.45 × 6.45 μm. Imaging was performed with a 100x/1.46 M27 OIL DIC Plan-Apochromat objective controlled by Zen Black software. A compact light source HXP120V mercury lamp was used for ocular observation. Data acquisition for mNG-IFT74 motility experiments was conducted using the confocal microscope in super-resolution mode with an Airyscan detector. The pixel dwell time was kept below 1 ms, and the scan time was below 200 ms (depending on scan speed), with 16-bit depth and a zoom range of 8-11 (depending on the length of the imaged cilia). Images were taken as a time series, with one image every 200 ms, for a total of 150 images.

Data collection for the experiment measuring mKATE2-IFT74 transport events in proximal and distal parts of the cilium (Fig. 6F-O) was performed using the confocal microscope in super-resolution mode with the Airyscan detector. The pixel dwell time was kept below 1 ms, and the scan time was below 200 ms or around 1 s, with 16-bit depth and a zoom range of 8-11. Images were taken as a time series, with one image every 200 ms or 1 s, for a total of 150 images. A 50 or 60% power of the 561 nm solid state laser was used to bleach the accumulated mKATE2-tagged IFT74 at the primary cilia tip.

The analysis of primary cilia length, transport frequency, and velocity was conducted in ImageJ. Cilia length was measured by tracing the mNG-IFT74 fluorescent signal with a segmented line. The frequency and velocity of IFT74 were determined using Multi Kymograph analysis in ImageJ, as previously described (30). Briefly, frequency corresponds to the number of anterograde or retrograde trajectories observed per second (Hz). The velocity of individual IFT trains was determined by multiplying the tangent of the angle (between each measured trajectory and the horizontal axis) of observed traces by pixel size and frames per second. To obtain frequency and velocity values, we only tracked trajectories with clear signs of directionality (“zig-zag” trajectories corresponding to the diffusive motion were not included in the analyses), that were easily distinguishable from the background signal. To measure the proximal and distal velocity of mKATE2-IFT74 (Fig. 6I-J, N), the trajectory angle of each mKATE2-IFT74 train was measured in the proximal and distal regions of the cilium.

### Time-lapse live cell imaging using TIRF microscopy

Wide-field techniques like TIRF microscopy offer more effective imaging and less phototoxicity compared to confocal microscopy, pertinent to studying IFT train trajectories. As primary cilia are typically positioned outside of the ∼300 nm illumination depth of true TIRF microscopy, we operated the microscope in VAEM mode, also known as pseudo TIRF (39). Time-lapse live cell imaging of all variants of mNG-KIF14 (with or without co-expression of mKATE2-IFT74) and mNG-IFT74-positive primary cilia was performed using live-cell TIRF microscopy (Fig. 3A-C, F, Suppl. Fig. 3C-F, Fig. 4D-E, Suppl. Fig. 4E-H, Fig. 6A-E, Suppl. Fig. 6C). To achieve live-cell TIRF, we used sub-critical angle incident light, which can illuminate fluorophores deeper in the sample while still significantly improving the signal-to-noise ratio (30). Imaging was conducted on cells in Nunc™ glass-bottom dishes 24 hours after initiating serum starvation. The time-lapse imaging was carried out on an inverted widefield microscope Nikon Eclipse Ti-E, equipped with a piezo Z-stage, motorized XY stage, Perfect Focus System, automatized H-TIRF module, module for environmental control Okolab, Nikon CFI HP Apo TIRF 100x Oil objective, NA 1.49, EM CCD Andor iXon Ultra DU897, 512 × 512 pixels, pixel size 16 × 16 µm, and W-VIEW GEMINI image splitter, controlled by NIS-Elements software (version 5.21). Images were taken as a time series, with one image every 100 ms, for a total of 300-600 images.

In addition, the analysis of mKATE2-IFT74 transport events in proximal and distal parts of the cilium (Suppl. Fig. 6C) was performed using an inverted fluorescence widefield microscope based on Nikon ECLIPSE Ti2 microscope body and iLas 2 illumination device allowing ring-TIRF microscopy, motorized XY stage, Perfect Focus System, module for environmental control Okolab, Nikon Apo TIRF 60x Oil (NA 1.49), TRF89901v2 ET -405/488/561/640 nm Laser Quad Band Set, PRIME BSI, 2048 × 2048 pixels, pixel size: 6.5 × 6.5 μm, controlled by software NIS-Elements. Images were taken as a time series, with one image every 100 ms, for a total of 300-600 images. 40% power of the 561 nm laser was used to bleach the accumulated mKATE2-tagged IFT74 at the primary cilia tip.

The analysis of KIF14 and IFT74 frequency and velocity, captured using live-cell TIRF, was conducted as described earlier in the “Time-lapse live cell imaging using confocal microscopy” section.

### Fluorescence recovery after photobleaching (FRAP)

For photobleaching experiments, cells were seeded on IBidi ibiTreat m-Slide 8-well chamber slide and starved using FBS-free FluoroBrite imaging DMEM. Cells were imaged on inverted Zeiss LSM880 laser scanning confocal microscope mounted on an Axio Observer.7 stand, equipped with an Airyscan detector and Monochromatic camera Axiocam MRm (with CCD sensor and pixel size 6.45 × 6.45 μm). Imaging was performed with a 100x/1.46 M27 OIL DIC Plan-Apochromat objective controlled by Zen Black software. A compact light source HXP120V mercury lamp was used for ocular observation.

For FRAP data measurements following the bleaching of cilium tip (Fig. 2D-E), the pixel dwell time was kept below 1 ms, and the scan time was below 200 ms (depending on scan speed), with 16-bit depth, 1388 × 1040 pixel resolution, and zoom range of 8-11 (depending on the length of the imaged cilia). Images were taken as a time series, with one image every 200 ms, for a total of 150 images. For bleaching mNG-tagged IFT74 in the primary cilia tip, 10% of the 488 nm argon laser power was applied after 5 scans. 15-20 cilia were analyzed per condition and experiment.

FRAP data following whole cilium bleaching (Suppl. Fig. 3A, Fig. 4B-C, 5B-C) were acquired with 16-bit depth, 512 × 512 pixel resolution, and zoom of 15. Images in Suppl. Fig. 3A were captured as a time series, with one image taken every 1.2 seconds, for a total of 30 images. Images in Fig. 4B-C, 5B-C were captured as a time series, with one image taken every 15 seconds, for a total of 12 images. Bleaching of mNG-tagged IFT74, ARL13B, and all KIF14 variants in the entire primary cilium was performed with 50% (Fig. 4B-C) or 80% (Suppl. Fig. 3A, Fig. 5B-C) of the 488 nm argon laser power after 1 scan. 3-31 cilia were analyzed per condition and experiment.

### Motility on GMPCPP-stabilized microtubules *in vitro*

The *in vitro* measurements of KIF14 motility (Suppl. Fig. 4I-K) were done using extracts from cells expressing fluorescently tagged KIF14, mixed with GMPCPP-stabilized microtubules, by adopting protocols described earlier (42,68,69). Specifically, hTERT RPE-1 cells expressing mNG-KIF14 (WT or 1-708) were harvested by centrifugation at 450 x g for 10 minutes at 4°C. The cell pellet was lysed in 200 µl of lysis buffer (50mM HEPES, 0.5M EGTA pH 8.0, 100mM MgCl_2_, 10% Triton X-100, Protease Inhibitor Cocktail, all obtained from Merck/Sigma-Aldrich) for 7 minutes on ice with occasional pipetting up and down. The cell lysate was sonicated and centrifuged at 21 000 x g for 50 minutes at 4°C. The supernatant was gently collected and aliquoted into PCR tubes. The PCR tubes containing the cell lysate were snap-frozen in liquid nitrogen and stored at -80°C.

Microtubules were prepared as described previously (70). The tubulin ratio used for microtubule preparation was 20% HiLyte 647 of labeled tubulin (Cytoskeleton Inc) to 80% of unlabeled tubulin, which was isolated for porcine brains as described previously (71).

*In vitro* microscopy assays to assess the behavior of mNG-KIF14 and of mNG-KIF14 1-708 were carried out in in-house made experimental chambers. Experimental chambers were constructed using two hydrophobized coverslips separated by stripes of parafilm as described previously (72). Chambers were functionalized with a solution of anti-β tubulin antibodies in 1 mg/ml of PBS and passivated with 10 mg/ml F127 solution in PBS for at least 1 hour. Each channel was flushed with 20 µl of motility buffer (BRB80 - 80 mM Pipes pH 6.8, 2 mM MgCl_2_, 1 mM EGTA; supplemented with 1 mM ATP, 10 mM DTT, 20 mM D-glucose, 0.5 mg/ml casein, 0.1% Tween 20, 0.22 mg/ml glucose oxidase, and 0.02 mg/ml catalase, followed by incubation with 5 µl of microtubule solution. Glucose oxidase and catalase were added immediately prior to the experiment. The ATP was purchased from Jena Bioscience, all the other chemicals were from Merck/Sigma-Aldrich.

After 10 seconds unbound microtubules were flushed with 20 µl of motility buffer. Finally, a channel was flushed with 20 µl of cell lysate diluted in motility buffer. Channel was observed for 13 minutes with an image acquisition every 30 seconds.

Imaging was performed using a Nikon Eclipse Ti2 mounted with sCMOS PRIME BSI camera (Teledyne Photometrics) and an oil immersion Nikon Apo TIRF 60x 1.49 N.A. objective. Fluorophore excitation was achieved with laser wavelengths of 640 nm, 561 nm or 488 nm. Emission filters EM700/75, EM610/75, and EM500-545 were used. Image analysis was done in ImageJ. Signal intensity of mNG-KIF14-708 was measured at the end of a microtubule and within the microtubule body region. Intensity values were corrected by background intensity adjacent to the microtubule.

### Motility of KIF14 on *Trypanosoma brucei* axonemes

6 × 10^7^ procyclic *Trypanosoma brucei* cells of the SmOxP927 strain (73) were washed with PBS and lyzed in 1ml of PEME (100 mM PIPES pH 6.9, 1 mM MgSO_4_, 2 mM EGTA, 0.1 mM EDTA) with 0.5% Igepal CA-630. The chemicals were purchased from Merck/Sigma-Aldrich. The resulting insoluble fraction (cytoskeletons) was pelleted by centrifugation at 2,000 x g and resuspended in 0.1 ml of PEME. The cytoskeletons were further 10x diluted in BRB80, introduced into microscopy channels, and allowed to adhere to glass for 10 minutes. The surface of the channels was subsequently passivated by a 1 hour incubation with 1% Pluronic F-127 in PBS. Behavior of KIF14-eGFP, purified from SF9 cells (22), on the axonemes was imaged in experiment chambers similar to those described above. Instead of antibody functionalization dry channels were perfused with a solution of the cytoskeletons and these let to adhere to the surface for 10 minutes. Channel was then passivated with 10 mg/ml F127 solution in PBS for at least 1 hour. Finally, channel was perfused with 20 µl motility buffer and 20 µl of purified KIF14-eGFP followed by imaging. A channel was imaged every 0.5 seconds for 5 minutes.

### Western blot

Cells were lysed in SDS lysis buffer (50 mM Tris-HCl pH 6.8, 10% glycerol, 1% SDS, 0.01% bromphenol blue, 2.5% 2-mercaptoethanol, Protease Inhibitor Cocktail), lysates loaded onto 6-10% SDS-polyacrylamide gels, electrophoresed, and blotted onto polyvinylidene difluoride membranes (Merck/Millipore). The membranes were blocked with 5% nonfat milk in PBS-Tween buffer (10 mM Tris-HCl pH 7.4, 100 mM NaCl, and 0.05% Tween) and then incubated with primary antibodies diluted in blocking solution overnight at 4°C. After washing with PBS-Tween, the membranes were incubated with HRP-linked secondary antibodies for 1 hour at RT. After washing with PBS-Tween, chemiluminescent detection using ECL Prime Western Blotting Detection Reagent (GE Healthcare) was performed according to the manufacturer’s instructions. Chemiluminescent signal was revealed by ChemiDoc Imaging System (BioRad). Subsequent analyses of tiff images were done using ImageJ software. The primary and secondary antibodies used are listed in Table 1.

### Protein structure prediction using AlphaFold

Protein structures were predicted using AlphaFold (version 2.1.1) (74), utilizing the UniProt and PDB protein databases. The predicted structures were refined and visualized in ChimeraX (version 1.8).

### Statistical analysis

All statistical analyses were performed using GraphPad Prism. Data are presented as mean ± SD, or as mean ± SEM. Where possible, the graphs depict the individual data points (signal intensity, cilium length, velocity per IFT track, etc.) determined for each condition across the biological replicates. The corresponding sample size (N), and the character of depicted data points are shown in the figure legends. The experiments were typically conducted in triplicates (n = number of biological replicates) unless specified otherwise in figure legends. Statistical differences between groups were assessed using a t-test or ANOVA followed by Tukey’s multiple comparisons test. For all statistical analyses, P < 0.05 was considered significant (*P < 0.05, **P < 0.01, ***P < 0.001, and ****P < 0.0001).

## Acknowledgments

We thank Erich Nigg, Peter Jackson, and Francis Barr for providing reagents, Andrea Lacigová for help with stable cell lines preparation, David Vysloužil for assistance with AlphaFold analysis and cloning, and Tomáš Loja for assistance with cell sorting. The work was supported by a grant from the Czech Science Foundation (21-21612S) to LČ and MH. The work carried out in the laboratory of VV was supported by the INTER-ACTION grant LUAUS24182 by the Czech Ministry of Education, Youth and Sports. We acknowledge the core facility CELLIM and BIOCEV, both supported by the Czech-BioImaging large RI project (LM2018129 funded by MEYS CR) for their support with obtaining scientific data presented in this article.

## Authors contribution

EM - performed and analyzed most of the experiments and assembled figures; PP - performed and/or analyzed experiments in Fig. 1 and Suppl. Fig. 1, 6A-B, D-E; RP - performed and analyzed *in vitro* TIRF experiments; LŠ - contributed to optimization and analysis of live-cell TIRF experiments; MH - generated several reporter cell lines; VV - optimized use of *Trypanosoma* cytoskeletons in *in vitro* experiments, and provided reagents; ZL - supervised *in vitro* TIRF experiments; LČ - conceptualized and supervised the study, and together with EM wrote the paper. All authors commented on and approved the final version of the manuscript.

**Video S1**.

Bi-directional movement of mNG-IFT74 particles, corresponding to the anterograde and retrograde transport in mNG-IFT74 hTERT-RPE-1 reporter cell.

**Video S2**.

Bi-directional movement of mNG-IFT74 particles, corresponding to the anterograde and retrograde transport in mNG-IFT74 hTERT-RPE-1 KIF14-depleted reporter cell.

**Video S3**.

Motion of purified KIF14-eGFP along the axonemal microtubules in vitro, using single-molecule TIRF microscopy.

