## Supplemental Figures for "Kinesin-3 KIF14 Regulates Intraflagellar Transport Dynamics in Primary Cilia"

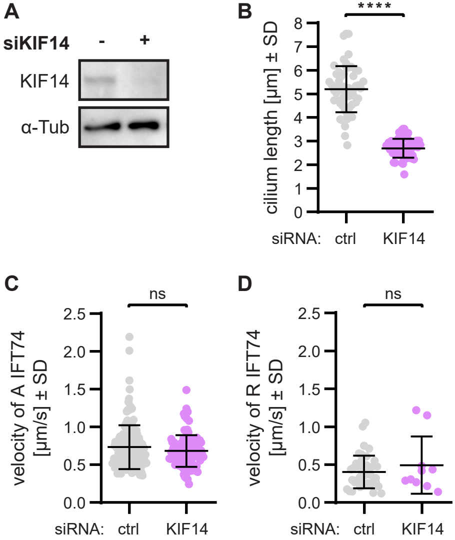


**Supplementary figure 1.**
**(A)** Western blot analysis of KIF14 siRNA efficiency. Alpha-tubulin (α-Tub) was used as a loading control. **(B)** Quantification of the length of mNG-IFT74 positive primary cilia following KIF14 depletion (unpaired t-test, ****$P < 0.0001$); n = 4, N = 57-58 (depicted data points represent values from individual cilia). **(C-D)** Quantification of mNG-IFT74 transport events shows no major difference in the velocity of anterograde **(C)** and retrograde **(D)** trains after KIF14 depletion (unpaired t-test); n = 4, N = 106-152 for C and 10-42 for D (depicted data points represent values from individual tracks).

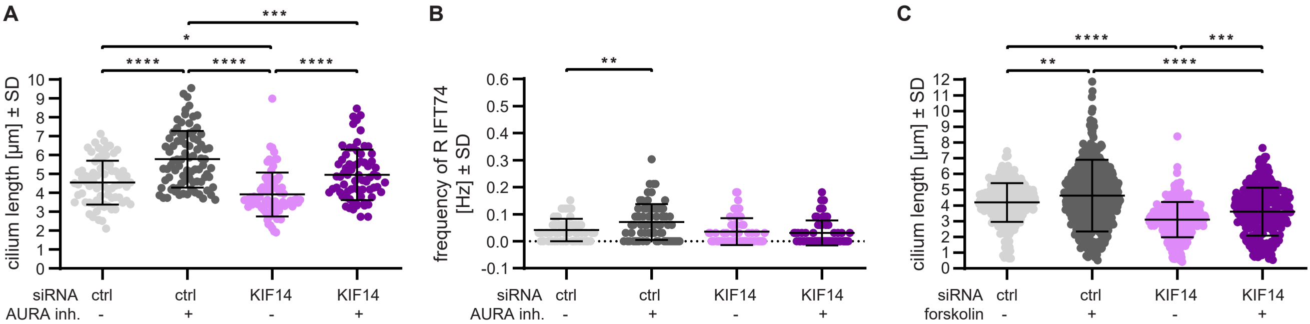


**Supplementary figure 2.**

**(A)** Quantification of the length of mNG-IFT74 positive primary cilia following KIF14 depletion and/or AURA inhibition (one-way ANOVA, $*P < 0.05$, $***P < 0.001$, $****P < 0.0001$); n = 3, N = 71-83, (depicted data points represent values from individual cilia). **(B)** Quantification of mNG-IFT74 transport events shows a lack of rescue of AURA inhibition of KIF14 depletion-mediated retrograde trains frequency reduction (one-way ANOVA, $**P < 0.01$); n = 3, N = 60-80 (depicted data points represent values from individual cilia). **(C)** Quantification of the length of primary cilia following KIF14 depletion and/or forskolin treatment (one-way ANOVA, $**P < 0.01$, $****P < 0.0001$); n = 2, N = 260-440 (depicted data points represent values from individual cilia).

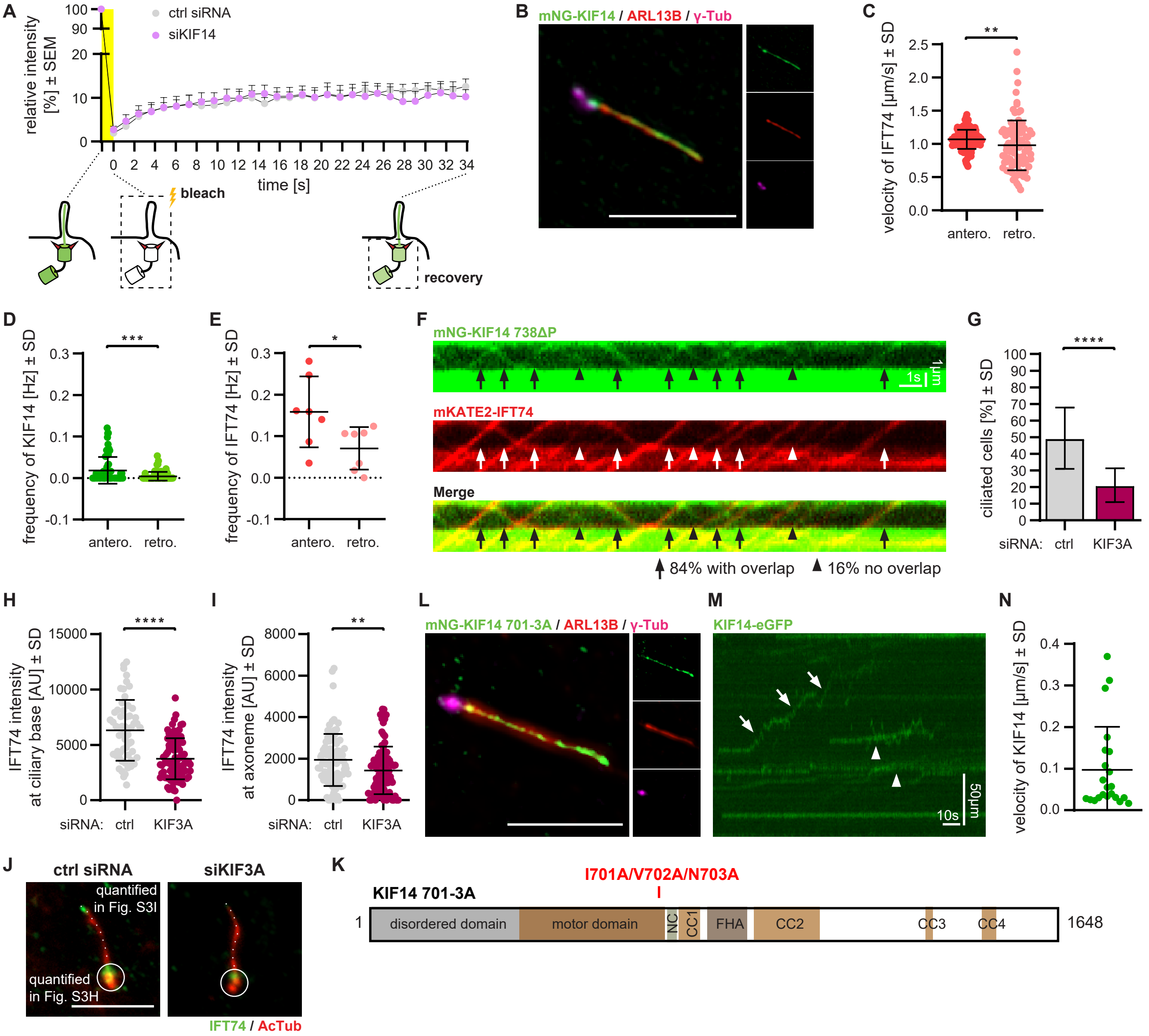


**Supplementary figure 3.**
**(A)** Measurement of recovery of mNG-IFT74 at the cilia base after KIF14 depletion following whole cilium bleach (see the scheme below the graph) using FRAP; n = 2, N = 21-31. **(B)** Localization of mNG-KIF14 in a primary cilium of hTERT RPE-1 cells by IF microscopy. ARL13B staining was used to detect primary cilia, and γ-Tub staining indicates centrioles; scale bar = 5 µm. **(C-E)** Quantification of anterograde and retrograde transport events (velocities and frequencies) determined using live-cell TIRF microscopy and kymograph analysis. Velocities of mNG-IFT74 trains are shown in **(C)** (unpaired t-test, $**P < 0.01$); n = 1, N = 95-174 (depicted data points represent values from individual tracks). The frequency of mNG-KIF14 moving particles is shown in **(D)** (unpaired t-test, $***P < 0.001$); n = 1, N = 67 (depicted data points represent values from individual cilia) **(E)** Frequency of mNG-IFT74 anterograde and retrograde trains, respectively (unpaired t-test, $*P < 0.05$); n = 1, N = 7 (depicted data points represent values from individual cilia). **(F)** Visualization of mNG-KIF14 738ΔP and mKATE2-IFT74 movement in the cilium of hTERT-RPE-1 cells using kymograph analysis. 84% of processive mNG-KIF14 738ΔP trajectories overlapped with those of mKATE2-IFT74 (arrows), while 16% of mNG-KIF14 738ΔP processive trajectories lacked corresponding mKATE2-IFT74 signal (arrowheads); scale bar = 1 µm and 5 s, respectively, n = 1, N = 31. **(G)** Quantification of the percentage of ciliated cells; n = 2, N = 496-508. **(H-I)** Quantification of the fluorescence intensity of IFT74 at the ciliary base **(H)** and along axoneme **(I)** after KIF3A depletion (unpaired t-test, $**P < 0.01$, $****P < 0.0001$); n = 2, N = 85-87, (depicted data points represent values of individual cilia analyzed). **(J)** Localization of IFT74 in a primary cilium of hTERT RPE-1 cells after KIF3A depletion. AcTub staining was used to detect primary cilia. The circles illustrate the cilium base, which were analyzed in H, the dashed lines illustrate the axoneme, which were analyzed in I; scale bar = 5 µm. **(K)** Schematic visualization of KIF14 functional domains and the location 701-3A mutation within KIF14 protein. NC - neck coil, CC - coiled coil, FHA - forkhead associated domain. **(L)** Localization of mNG-KIF14 701-3A mutant (I701A/V702A/N703A) in a primary cilium of hTERT RPE-1 cells detected by IF microscopy. ARL13B staining was used to detect primary cilia, and γ-Tub staining indicates centrioles; scale bar = 5 µm. **(M)** Visualization of KIF14-eGFP diffusive (arrowheads) and processive (arrows) movement along *Trypanosoma brucei* axonemal microtubules in vitro using kymograph analysis; scale bar = 1 µm and 5 s, respectively. **(N)** Quantification of KIF14-eGFP velocities in *Trypanosoma brucei*.

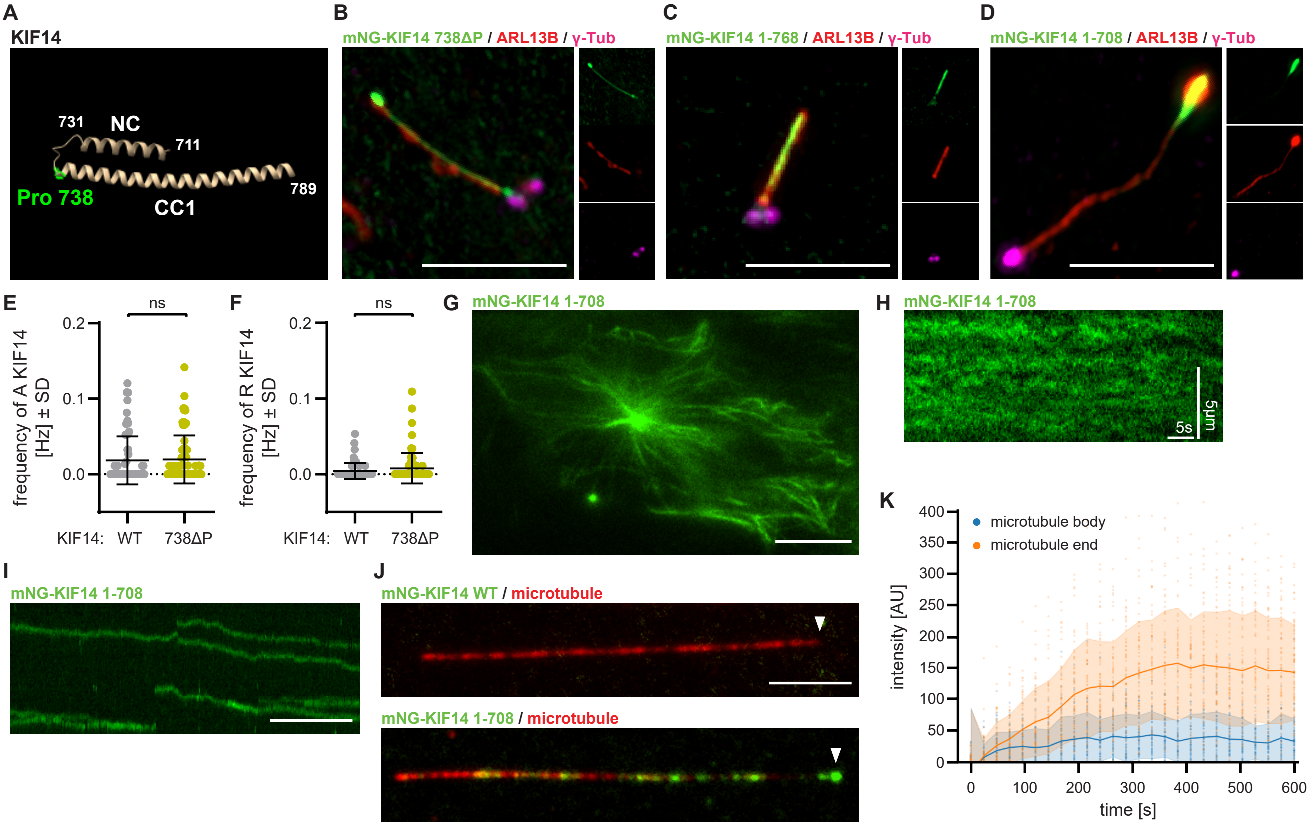


**Supplementary figure 4.**

**(A)** Identification of NC (711-731) and CC1 (738-789) domains and the first Proline (Pro 738) of CC1 domain based on AlphaFold structure prediction. (B-D) Localization of depicted mNG-tagged variants of KIF14 in a primary cilium of hTERT RPE-1 cells observed by IF microscopy. ARL13B staining was used to visualize primary cilia, γ-Tub staining labeled centrioles; scale bar = 5 µm. mNG-KIF14 738ΔP **(B)**, mNG-KIF14 1-768 **(C)**, and mNG-KIF14 1-708 **(D)**. **(E-F)** Quantification of the frequency of anterograde **(E)** and retrograde **(F)** processive runs of mNG-KIF14 and mNG-KIF14 738ΔP, determined using live-cell TIRF microscopy and kymograph analysis (unpaired t-test); n = 1, N = 68-71 (depicted data points represent values from individual tracks). **(G)** mNG-KIF14 1-708 localization to cytosolic microtubules. **(H)** Visualization of the diffusive motion of mNG-KIF14 1-708 along cytosolic microtubules using kymograph analysis; scale bar = 5 µm and 5 s, respectively. **(I)** Visualization of the processive motion of mNG-KIF14 1-708 on GMPCPP-stabilized microtubules using kymograph analysis; scale bar = 5 µm. **(J)** Representative snapshots from TIRF microscopy showing accumulation of mNG-KIF14 1-708, but not mNG-KIF14, at the ends of GMPCPP-stabilized microtubules; scale bar = 5 µm. Quantification of the signal of mNG-KIF14 1-708 at the body versus the end of GMPCPP-stabilized microtubules, measured by TIRF, is shown in **(K)**; n = 1, N = 69.

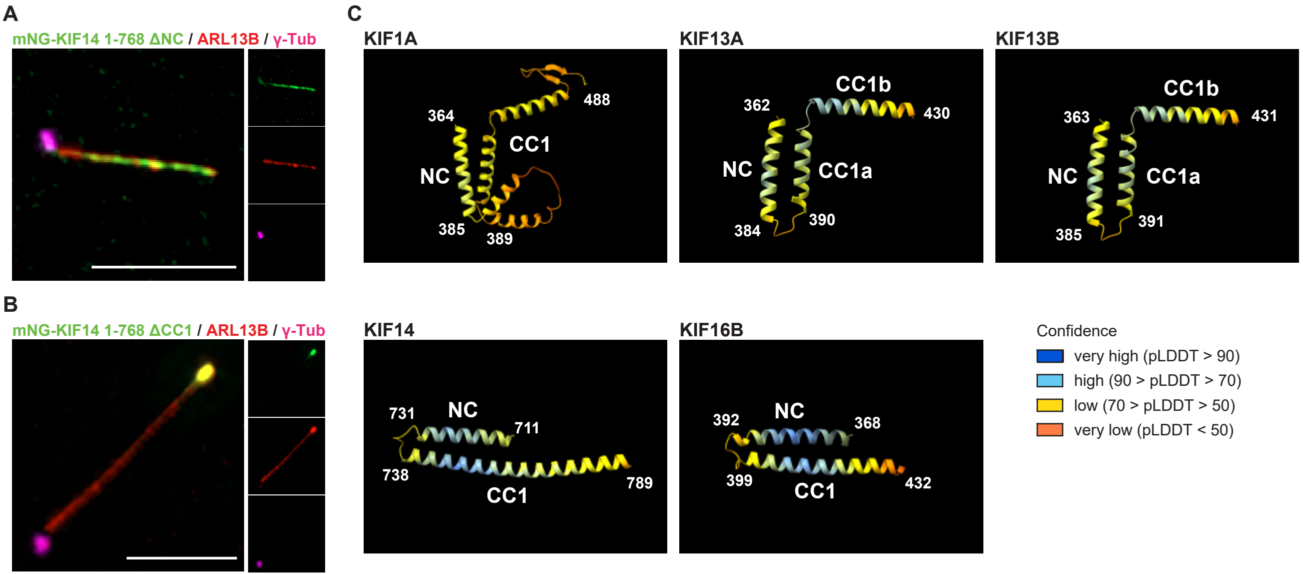


**Supplementary figure 5.**
Localization of mNG-KIF14 1-768 ΔNC **(A)**, and mNG-KIF14 1-768 ΔCC1 **(B)** in the cilium of hTERT RPE-1 cells, detected by IF microscopy. ARL13B staining was used to visualize primary cilia, and γ-Tub staining labeled centrioles; scale bar = 5 µm. **(C)** Identification of NC and CC1 domains for KIF1A, KIF13A, KIF13B, KIF14, and KIF16B based on AlphaFold structure prediction. The color code illustrates the confidence of Alphafold predictions based on pLDDT score (per-residue measure of local confidence); pLDDT ≥ 90 (very high confidence, dark blue), pLDDT 70-90 (high confidence, light blue), pLDDT  50-70 (low confidence, yellow), and pLDDT < 50 (very low confidence, red).

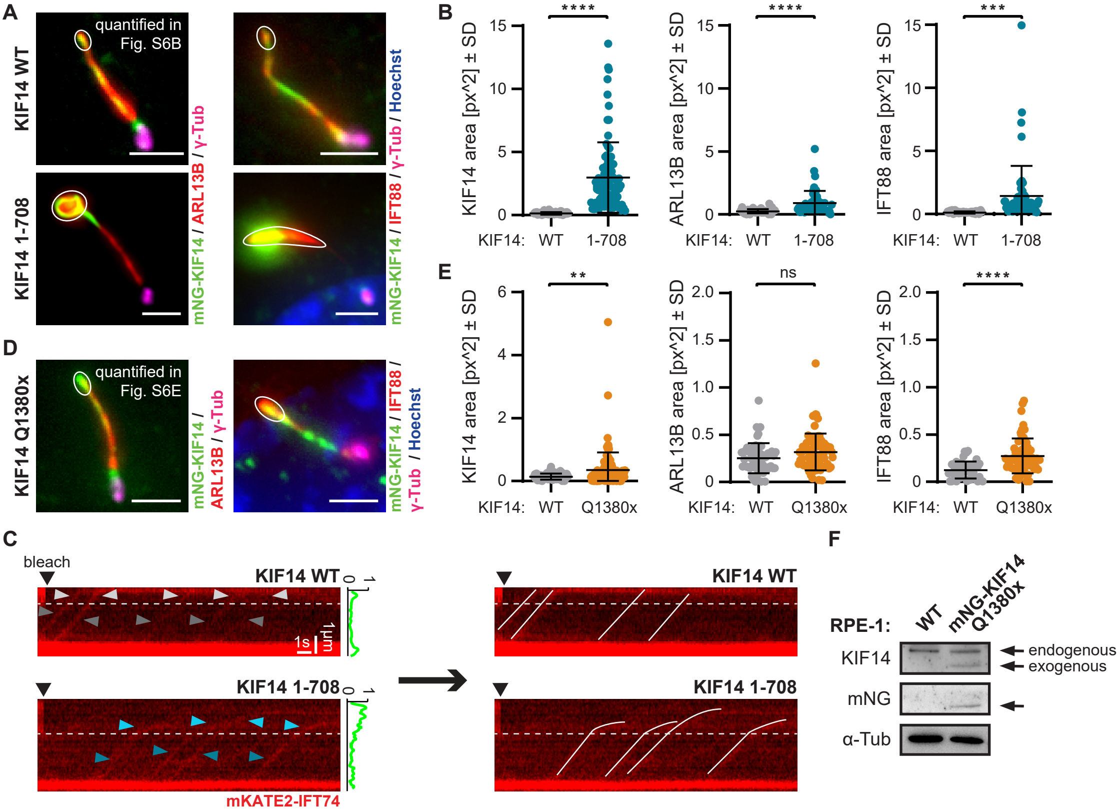


**Supplementary figure 6.**

**(A)** Distribution of KIF14, ARL13B, and IFT88 in the presence of mNG-KIF14 WT or mNG-KIF14 1-708 in primary cilia of hTERT RPE-1 cells by IF microscopy. Note the accumulation of KIF14/ARL13B/IFT88 in the distal part of the cilium in mNG-KIF14 1-708 condition. The white circles illustrate the distal part of the cilium, which was analyzed in B; scale bar = 2 µm. **(B)** Analysis of the area with the visible signal of KIF14, ARL13B, or IFT88 in the distal part of cilia in the presence of KIF14 WT or KIF14 1-708 (unpaired t-test, ***P < 0.001, ****P < 0.0001); n = 3, N = 46-83, (depicted data points represent values of individual cilia analyzed). **(C)** Kymograph visualization showing mKATE-IFT74 slowing down in the distal part of cilium containing high density of KIF14 1-708 (dark blue arrowheads for proximal part and light blue arrowheads for distal part of primary cilium) compared to KIF14 WT (dark gray arrowheads for proximal part and light gray arrowheads for distal part of primary cilium) using live-cell TIRF; scale bar = 1 µm and 1 s, respectively. The plot profile next to the kymograph illustrates the distribution of mNG-KIF14 WT relative to mNG-KIF14 1-708 along the primary cilium. **(D)** Distribution of KIF14, ARL13B, and IFT88 in primary cilia in the presence of mNG-KIF14 Q1380x in hTERT RPE-1 cells, observed by IF microscopy. Note the accumulation of KIF14/IFT88 in the distal part of the cilium in mNG-KIF14 Q1380x condition. The white circles highlight the distal part of the cilium, which was analyzed in E; scale bar = 2 µm. **(E)** Analysis of the area with the visible signal of KIF14, IFT88, or ARL13B in the distal part cilia in the presence of KIF14 Q1380x (unpaired t-test, **P < 0.01, ****P < 0.0001); n = 3, N = 46-118, (depicted data points represent values of individual cilia analyzed). **(F)** Western blot analysis of KIF14 Q1380x level. Alpha-tubulin (α-Tub) was used as a loading control.
