## Supplementary material for "Kinesin-3 KIF14 Regulates Intraflagellar Transport Dynamics in Primary Cilia": Table 1

| REAGENT or RESOURCE | SOURCE | IDENTIFIER |
|---|---|---|
| **Antibodies** | | |
| Rabbit polyclonal Anti-ARL13B | Proteintech | 17711-1-AP |
| Mouse monoclonal Anti-γ-Tubulin | Sigma-Aldrich/Merck | T6557 |
| Mouse monoclonal Anti-Acetylated-Tubulin | Santa Cruz | sc-23950 |
| Rabbit polyclonal Anti-IFT74 | Proteintech | 27334-1-AP |
| Rabbit polyclonal Anti-IFT88 | Proteintech | 13967-1-AP |
| Rabbit polyclonal Anti-KIF14 | Abcam | ab3746 |
| Rabbit monoclonal Anti-mNeonGreen Tag | Cell Signaling | 55074S |
| Mouse monoclonal Anti-α-Tubulin | Sigma-Aldrich/Merck | T9026 |
| Mouse monoclonal Anti-β-Tubulin I | Sigma-Aldrich/Merck | T7816 |
| Donkey anti-Rabbit IgG, Alexa Fluor 488 | Thermo Fisher Scientific | A21206 |
| Donkey anti-Mouse IgG, Alexa Fluor 568 | Thermo Fisher Scientific | |
| Donkey anti-Rabbit IgG, Alexa Fluor 568 | Thermo Fisher Scientific | A10042 |
| Donkey anti-Mouse IgG, Alexa Fluor 647 | Thermo Fisher Scientific | A31571 |
| Goat anti-Mouse IgG, HRP-linked | Sigma-Aldrich/Merck | A4416 |
| Goat anti-Rabbit IgG, HRP-linked | Cell Signaling | 7074S |
| **Chemicals** | | |
| Acrylamide | Serva | 1068701 |
| ATP | Jena Bioscience | NU-1010 |
| BSA | Sigma-Aldrich/Merck | A9647 |
| β-Casein | Sigma-Aldrich/Merck | C6905 |
| Catalase | Sigma-Aldrich/Merck | C40 |
| Clarity Western ECL Substrate | Bio-Rad | 1705061 |
| D-Glucose | Sigma-Aldrich/Merck | G7528 |
| DL-Dithiothreitol | Sigma-Aldrich/Merck | 43819 |
| DMEM, high glucose, GlutaMAX™ Supplement, pyruvate | Gibco/ThermoFisher | 31966021 |
| DMEM/F-12, no glutamine | Gibco/ThermoFisher | 21331020 |
| EGTA | Sigma-Aldrich/Merck | E4378 |
| FBS Ultra-low endotoxine | Biosera | FB-1101/500 |
| FluoroBrite DMEM | Thermo Fisher | A1896701 |
| Glucose Oxidase | Sigma-Aldrich/Merck | G2133 |
| L-glutamine | Biosera | XC-T1715/100 |
| IGEPAL CA-630 | Sigma-Aldrich/Merck | I3021 |
| Lipofectamine™ RNAiMAX | Invitrogen | 13778150 |
| 2-mercaptoethanol | Sigma-Aldrich/Merck | M3148 |
| $MgCl_2$ | Sigma-Aldrich/Merck | M2670 |
| NaCl | Penta/Merci | 7647-14-5 |
| OPTIMEM | Gibco/ThermoFisher | 31985070 |
| Penicillin/Streptomycine | Biosera | XC-A4122/100 |
| PIPES | Sigma-Aldrich/Merck | P6757 |
| Pluronic F-127 | Sigma-Aldrich/Merck | P2443 |
| Polybrene | Sigma-Aldrich/Merck | H9268 |
| Polyethylenimine (PEI) | Sigma Aldrich/Merck | 408727 |
| ProLong™ Glass Antifade Mountant | Invitrogen/ThermoFisher | P36980 |
| cOmplete™, Mini, EDTA-free Protease Inhibitor Cocktail | Roche/Merck | 11836170001 |
| SDS | Supelco/Merck | 74255 |
| TCS7010 | Tocris Bioscience | 5286 |
| Triton X-100 | Sigma-Aldrich/Merck | X100 |

| | | | |
|---|---|---|---|
| Trizma® base | | Sigma-Aldrich/Merck | T1503 |
| Tween 20 | | Sigma-Aldrich/Merck | P1379 |
| **Other** | | | |
| Gateway™ LR Clonase™ II Enzyme mix | | Invitrogen/ThermoFisher | 11791020 |
| glass bottom dish with 20 mm micro-well | | Cellvis | D35-20-1.5-N |
| HiLyte Fluor™ 647 labeled microtubules | | Cytoskeleton | TL670M |
| PES syringe filters 0.45 µm | | filtraTECH | SF33PE45S |
| Immobilon® -P PVDF Membrane | | Millipore/Merck | IPVH00010 |
| QuikChange XL Site-Directed Mutagenesis Kit | | Agilent Technologies | 50-125-162 |
| µ-Slide 8 well chamber slide | | IBidi | ibiTreat/80826 |
| Zero Blunt™ TOPO™ PCR Cloning Kit | | Invitrogen/ThermoFisher | 450245 |
| Zeiss Immersion oil Immersiol 518 F | | Carl Zeiss | 444960-0000-000 |
| **Experimental models: Cell lines** | | | |
| hTERT-RPE-1 | | a gift from E. Nigg | (75) |
| hTERT-RPE-1 Flp-In™ T-REx™ | | a gift from E. Nigg | (76) |
| Phoenix-Ampho | | ATCC | CRL-3213 |
| **Primers (sequence 5´->3´)** | | | |
| KIF14 738ΔP forward | agtcggaatattgacgaacgatacaggctc | | |
| KIF14 738ΔP reverse | gagcctgtatcgttcgtcaatattccgact | | |
| KIF14 1-768 reverse | tca cactctttgcatttctgccat | | |
| KIF14 1-708 reverse | tca atttactttagcaatgttga | | |
| KIF14 1-768 ΔNC forward | ccgtttaatagtcaacattgctaaagtaaatattgaccctgaacgata | | |
| KIF14 1-768 ΔNC reverse | tatcgttcagggtcaatatttactttagcaatgttgactattaaacgg | | |
| KIF14 1-768 ΔCC1 forward | tgctcagagaaacagtcggaattgaaagggtggg | | |
| KIF14 1-768 ΔCC1 reverse | cccaccctttcaattccgactgtttctctgagca | | |
| KIF14 701-3A forward | gcgttcatatcttcatttactttagcaatggcggctgctaaacgggcttggttagcatatctaagtgtgc | | |
| KIF14 701-3A reverse | gcacacttagatatgctaaccaagcccgtttagcagccgccattgctaaagtaaatgaagatatgaacgc | | |
| KIF14 Q1380x forward | tgctaactgccccacatactttacctattgttgtacaatttggattgcattctttt | | |
| KIF14 Q1380x reverse | aaaagaatgcaatccaaattgtacaacaataggtaaagtatgtggggcagttagca | | |
| **siRNA (targeted sequence (5'->3'))** | | | |
| Control siRNA | Silencer® Firefly Luciferase (GL2 + GL3) | Thermo Fisher Scientific | AM4629 |
| KIF14 | #1: TGACACGAAACAAGTTTATA | Thermo Fisher Scientific | s19265 |
| | #2: TTTAATAGTCAACATTGCTA | Thermo Fisher Scientific | s19263 |
| KIF3A | - | Thermo Fisher Scientific | s21942 |
| **Software and algorithms** | | | |
| AlphaFold | | (74) | version 2.1.1 |
| ChimeraX | | (77) | version 1.8 |
| GraphPad Prism | | GraphPad Software | version 8.0.1 |
| ImageJ | | (67) | version 1.54k |
| ZEN Black | | Carl Zeiss | version 2.3 SP1 |
| ZEN Blue | | Carl Zeiss | version 2.6 |
| NIS-Elements | | Nicon Instruments Inc. | version 5.21 |
